# Vertical RAS-pathway inhibition in pancreatic cancer drives therapeutically exploitable mitochondrial alterations

**DOI:** 10.1101/2025.02.03.636222

**Authors:** Philipp Hafner, Steffen J. Keller, Xun Chen, Asma Alrawashdeh, Huda Jumaa, Friederike I. Nollmann, Solène Besson, Judith Kemming, Oliver Gorka, Tonmoy Das, Bismark Appiah, Ariane Lehmann, Mujia Li, Petya Apostolova, Bertram Bengsch, Stefan Tholen, Oliver Schilling, Olaf Groß, Andreas Vlachos, Uwe A. Wittel, Dominik von Elverfeldt, Wilfried Reichardt, Melanie Boerries, Geoffroy Andrieux, Guus J. Heynen, Stefan Fichtner-Feigl, Luciana Hannibal, Dietrich A. Ruess

**Affiliations:** Department of General and Visceral Surgery, Center for Surgery, Medical Center – University of Freiburg, Faculty of Medicine, Freiburg, Germany; University of Freiburg, Faculty of Biology, Freiburg, Germany; Department of Hepatobiliary and Pancreatic Surgery, The Affiliated Cancer Hospital of Zhengzhou University & Henan Cancer Hospital, Zhengzhou, 450008, China; Institute of Neuropathology, Medical Center – University of Freiburg, Faculty of Medicine, Freiburg, Germany; Institute of Medical Bioinformatics and Systems Medicine, Medical Center – University of Freiburg, Faculty of Medicine, Freiburg, Germany; Institute of Surgical Pathology, Medical Center – University of Freiburg, Faculty of Medicine, Freiburg, Germany; Division of Hematology, University Hospital Basel, Basel, Switzerland; Department of Internal Medicine II, Medical Center - University of Freiburg, Freiburg, Germany; CIBSS – Centre for Integrative Biological Signalling Studies, University of Freiburg, Freiburg, Germany; German Cancer Consortium (DKTK), Partner Site Freiburg, Freiburg, Germany, and German Cancer Research Center (DKFZ), Heidelberg, Germany; Department of Neuroanatomy– University of Freiburg, Faculty of Medicine, Freiburg, Germany; Division of Medical Physics, Department of Diagnostic and Interventional Radiology, University Medical Center Freiburg, Faculty of Medicine, University of Freiburg, Freiburg, Germany; Division of Hematology, Oncology and Tumor Immunology, Medical Department – Charité Universitätsmedizin Berlin, Berlin, Germany; Department of Pediatrics, Medical Center – University of Freiburg, Faculty of Medicine, Freiburg, Germany

**Keywords:** Pancreatic ductal adenocarcinoma, SHP2 inhibition, MEK1/2 inhibition, Mitochondrial alterations

## Abstract

**Background & Aims:** Oncogenic KRAS mutations drive metabolic rewiring in pancreatic ductal adenocarcinoma (PDAC). Src-homology 2 domain-containing phosphatase 2 (SHP2) is essential for full KRAS activity and promising dual SHP2/mitogen-activated protein kinase (MAPK) inhibition is currently being tested in clinical trials. Exploitable metabolic adaptations may contribute to an invariably evolving resistance.

**Methods:** To understand the metabolic changes induced by dual inhibition, we comprehensively tested cell lines, endogenous tumor models, and patient-derived organoids representing the full spectrum of PDAC molecular subtypes.

**Results:** We find that dual SHP2/mitogen-activated protein kinase kinase (MEK1/2) inhibition induces major mitochondrial alterations, elevates reactive oxygen species (ROS) levels and triggers a lipid peroxidase dependency. While anabolic pathways, glycolysis and autophagy were also affected, mitochondrial alterations persisted longterm into a therapy resistant state.

**Conclusions:** The resulting vulnerability to induction of ferroptotic cell death via combined SHP2/MEK1/2 and glutathione peroxidase (GPX4) inhibition provides a metabolic lever to reinforce RAS-pathway inhibition for targeted PDAC treatment.

## Introduction

Pancreatic ductal adenocarcinoma (PDAC) remains one of the deadliest cancers, with an estimated rise to the second leading cause of cancer-related deaths in the US within the coming years [1]. Despite recent therapeutic advances, PDAC still has a poor prognosis, with over 85% of patients succumbing to the disease within five years [2]. Surgery is the only curative option, but only about 20% of cases are resectable, and recurrence is common even after adjuvant chemotherapy [3]. The genetic landscape of PDAC has been extensively studied and it is clearly evident that oncogenic *KRAS* mutations are the dominant drivers of this entity, with more than 90% of PDAC tumors displaying activating alterations in the *KRAS* gene [4]. *KRAS* wild-type PDAC frequently harbor genetic alterations in other RAS pathway genes [5], confirming a high level of dependence of the entity on this signaling cascade. *KRAS* drives a variety of cellular functions including cell proliferation, differentiation, survival, and importantly, metabolic adaptation [6–11].

PDAC can be subclassified based on clinically relevant transcriptional expression profiles [12–15]. PDAC patients with tumors with a more dedifferentiated and mesenchymal expression pattern (basal-like/quasi-mesenchymal/squamous subtypes) appear to have a distinctly worse prognosis compared to patients with tumors with a differentiated, epithelial expression pattern (classical/progenitor subtypes) [12–16]. Initially, epithelial subtypes were found to correlate with KRAS dependency [17, 18], more recently an increase in gene dosage of mutated *KRAS* was described to directly impact PDAC biology and determine a more dedifferentiated and more aggressive phenotype [13, 19, 20].

Further, three metabolic subtypes of PDAC have been described, differing mainly in their dependence on glycolytic, lipogenic and redox pathways, with a suggested correlation with transcriptional signatures [21].

Previous studies have demonstrated that dual “vertical” RAS-pathway inhibition by targeting Src homology region 2 domain-containing phosphatase-2 (SHP2) and the mitogen-activated protein kinase kinase (MAP2K1/2) MEK1/2 or extracellular signaling kinases (MAPK3/1) ERK1/2 leads to synergistic growth inhibition of PDAC tumors [22, 23]. Clinical trials (e.g. NCT04916236) are currently underway to explore the potential of this approach for patients. Additionally, the rapidly expanding field of direct KRAS and RAS inhibition has already made tremendous impact [24–28]. However, as with MEK or ERK inhibitors, quickly evolving therapy resistance needs to be anticipated and combination strategies will likely be required to achieve durable responses. Co-targeting SHP2 bears promise in this context [29].

We here aimed at uncovering therapeutically exploitable PDAC-subtype dependent metabolic vulnerabilities evolving with indirect dual interference with RAS-pathway activity via vertically combined SHP2/MEK1/2 inhibition.

### Cell Culture Summary

The human pancreatic ductal adenocarcinoma (PDAC) cell lines MIA PaCa II, PANC-1, and YAPC were cultured in RPMI 1640 medium with 10% FCS and Penicillin-Streptomycin. Primary murine PDAC cells were derived from Kras^tm1Tyj^: *Kras^LSL-G12D/+^* + Trp53^tm1Brn^:*Trp53^fl/fl^* + Ptf1a^tm1(Cre)Hnak^: *Ptf1a^Cre-ex1/+^* (KPC) mice tumors and prepared into fragments for culture. Additionally, primary murine PDAC cell lines (F2453 and F2683) from *Kras^FSF-G12D/+^; Trp53^frt/frt^; Ptf1a-^Flpo/+^; R26-FSF-CreERT; Ptpn11^loxp/loxp^* (KPF) models, with prior Ptpn11 deletion via 4-OH-tamoxifen, were also used. All cells were maintained at 37°C, 5% CO2, and routinely tested for mycoplasma. For therapy resistant cell lines human and murine PDAC cells were cultured in RPMI 1640 with 10% FCS and Penicillin-Streptomycin, and treated continuously with SHP099, Trametinib, or both for 8 weeks, with media refreshed every three days.

### Human PDAC Organoid Generation

PDAC tissues were minced, collagenase-treated, and ACK-lysed to isolate cells, which were then cultured as organoids in Matrigel domes with human feeding medium, following protocols adapted from the Tuveson Lab [30, 31].

### Plasmids and Transfection

CRISPR-Cas9 constructs targeting the PTPN11 gene were designed with pX458 vectors containing specific guide RNAs (gRNA 1 and gRNA 2). The oligonucleotide sequences were as follows: gRNA 1 - forward: CACCGGAGGAACATGACATCGCGG, reverse: AAACCCGCGATGTCATGTTCCTCC; gRNA 2 - forward: CCACGAACATGACATCGCGGAGGTG, reverse: AAACCACCTCCGCGATGTCATGTTC. Each gRNA oligo pair was annealed and ligated into Bbs1-digested pX458 vectors. MIA PaCa II cells were transfected with pX458-PTPN11-gRNA plasmids using FuGENE® HD Transfection Reagent. Successfully transfected cells were sorted as singlets based on GFP expression and grown in a 96-well plate. SHP2-knockout clones, verified by Western Blot, were named by gRNA, e.g., MIA PaCa II KO2 for gRNA2.

### Mice

All experiments were conducted at the Universitätsklinikum Freiburg. The animals were kept in hygienic, pathogen-free conditions, and all animal experiments and care were in accordance with the law and with guidelines of institutional committees, and they were approved by the local authority (Regierungspräsidium AA; approval number G/19-137). The authors complied with the “Animal Research: Reporting of in Vivo Experiments” (ARRIVE) guidelines.

### Drugs and antibodies

The drugs SHP099 (S8278), Trametinib (S2673), RMC-4550 (S8718) and ML-210 (S0788) were purchased from Selleckchem. The metabolic related drugs Oligomycin A (11342), Rotenone (13995), Antimycin A (19433), 2-Deoxyglucose (14325) and 6-Aminonicotinamide (10009315) were purchased from Cayman Chemical. Chloroquine (HY-17589A), PGC-1α-inhibitor (HY-101491) and C75 (HY-12364) were purchased from MedChemExpress. Etomoxir (E1905) and *tert*-Butyl hydroperoxide (TBHP) (458139) were obtained from Sigma-Aldrich and carbonyl cyanide-p-trifluoromethoxyphenylhydrazone (FCCP) (ab147482) from Abcam. Proteins were detected by western blot with the following antibodies: pSTAT3 1:1000 (D3A7), STAT3 1:1000 (79D7), SHP2 1:1000 (D50F2), pAKTS473 1:1000 (D9E), AKT 1:1000 (C67E7), pERKT202/Y204 1:1000 (D13.14.E4), ERK 1:1000 (137F5), KRAS 1:1000 (E2M9G), KRASG12D 1:1000 (D8H7), KRT81 1:1000 (H00003887-M01J), ZEB1 1:1000 (D80D3), E-Cadherin 1:1000 (24E10), N-Cadherin 1:1000 (D4R1H), Vimentin 1:1000 (D21H3), GATA-6 1:1000 (D61E4), SLUG 1:1000 (C19G7), SNAIL 1:1000 (C15D3) and Actin 1:5000 (A2066). All antibodies were purchased from Cell Signaling Technologies except KRT81 (Thermo Fisher Scientific) and Actin (Sigma-Aldrich).

### *In vitro* growth assay

For this cell proliferation assay, we cultured the following PDAC cells at the corresponding densities: murine KPC cells at 3.125 cells/cm^2^ surface area, human PANC-1 and MIA PaCa-II cells at 5.800 cells/cm^2^ surface area, and human YAPC cells at 7.800 cells/cm^2^ surface area. The cells were plated overnight to allow attachment to the surface area. After 12-24 hours, we added either 15 μM of SHP099, 10 nM of Trametinib, or a combination of both as targeted therapy. Additionally, we added metabolic inhibitors at the indicated concentrations. The experiment was terminated when the cells with the respective targeted therapy reached a cell density of approximately 90%. At the end of the experiment, cells were fixed with 3,8% formaldehyde and stained with a solution of crystal violet (20% Thanol+0.2% crystal violet from Sigma-Aldrich). Fixed and stained cells were imaged with ChemiDoc^TM^ MP Imaging System from Bio-Rad and mean grey value was determined using ImageJ version 1.53a.

### *In vitro* growth assay of Human PDAC organoids

A total of 1,000 cells were mixed with 5 μl Matrigel, and one dome was placed in each well of a 96-well plate. The cells were cultured in human feeding medium for 72 hours. On the following day, cells were treated with either 15 μM SHP099, 25 nM Trametinib, or a combination of both. Additionally, organoids were treated with metabolism-associated inhibitors at the indicated concentrations. After 5 days of treatment, organoid proliferation was quantified using the BrdU colorimetric immunoassay from Roche (Cat. No. 11 647 229 001).

### Flow Cytometry

Human and murine PDAC cells were plated and allowed to attach overnight following treatment with 15 μM SHP099, 10 nM Trametinib, or both. After 72 hours, intracellular ROS levels were assessed using a BD LSRFortessa flow cytometer with a 20 nm bandpass at 712 nm. To induce ROS, the cells were treated with 100 μM tert-butyl hydroperoxide (TBHP) 30 minutes before harvesting. TBHP treated cells were used as a positive control. For mitochondrial mass, cells were stained with MitoTrackerTM Deep Red FM following similar steps.

### Immunoblotting

Cells were lysed with RIPA buffer and protein concentrations determined. Lysates were denatured, loaded onto SDS-PAGE, and transferred to nitrocellulose. After blocking, membranes were incubated with primary and secondary HRP-conjugated antibodies, then visualized with ChemiDoc.

### Oxygen and glucose consumption

To measure OCR and ECAR of PDAC cells, 3×10□ PANC-1 cells were plated in a Poly-D-Lysin pre-coated XF96 microplate and allowed to attach overnight. Cells were treated for 24 hours with 15 μM SHP099, 10 nM Trametinib, or their combination. On the day of measurement, the medium was replaced with DMEM containing glucose (25 mM) and glutamine (2 mM) without sodium bicarbonate (pH 7.4) and incubated for 30-60 minutes at 37°C in a CO_₂_-free incubator. Mitochondrial stress (Seahorse 101706-100) and glycolysis stress test (Seahorse 103020-100) was performed based on manufactures instructions and OCR and ECAR were normalized to cell number using the CyQUANT™ assay.

### Histology and immunohistochemistry

Tissue samples were fixed in 4% paraformaldehyde, dehydrated, and embedded in paraffin. Sections (4 μm) underwent deparaffinization and antigen retrieval with 10 mM citrate buffer (pH 6.0), followed by 3% hydrogen peroxide blocking. Core staining was done with Hemalaun. Immunohistochemistry used anti-MDA antibody (1:300 Abcam, Ab27642), and signal intensity was analyzed using QuPath (v0.2.2). To interpret the signal intensity of tumor cells a histoscore was calculated that consists of the sum of (1 x % weak stainig) + (2 x % moderate stainig) + (3 x % strong stainig).

### Transcriptomics

To analyze transcriptomic profiles cells were treated with 15 μM SHP099, 10 nM Trametinib, or their combination. After 48 hours, cells were lysed in QIAGEN’s RLT Plus buffer and purified using the RNeasy® Plus Mini Kit. Sequencing was conducted at DKFZ using Illumina’s NovaSeqTM6000. Data were converted to FASTQ files, quality-trimmed, and aligned to reference genomes (GRCh38 for human, GRCm39 for mouse) with STAR aligner (v2.7.10a) [32, 33]. Differential expression analysis was performed using limma, with GSEA carried out using fgsea and MSigDB collections [34–36]. PDAC subtypes were classified based on five key studies (Moffitt, Bailey, Collisson, Puleo, Chan-Seng-Yue), combining subtype signatures into Basal and Classical categories. Subtype classification per sample used single-sample gene set enrichment analysis (ssGSEA) via the clusterProfiler package [37]. Human PDAC signatures were mapped to mouse genes, and CIBERSORTx estimated cell-type composition in 20 mouse PDAC samples using a custom 13-cell-type reference matrix [38].

### Proteomics

Proteomic samples were prepared, TMT-labeled, and fractionated following Alatibi et al., 2021 [39]. For LC-MS/MS, 800 ng of peptides were analyzed on a Q-Exactive Plus mass spectrometer connected to an EASY-nLCTM 1000 UHPLC system. The column configuration included an Acclaim™ PepMap™ C18 column and a 200 cm µPac GEN1 analytical column, with peptide separation achieved using a 120-minute gradient from 5% to 100% buffer B (0.1% formic acid in 80% acetonitrile). Peptides were analyzed in data-dependent acquisition mode, with survey scans at 70,000 resolution and targeted fragmentation of the top 10 precursor ions at 17,500 resolution. Data were processed using MaxQuant v1.6.14.0 with the Human-EBI database (downloaded January 9, 2020) [39].

### Preparation of samples for metabolomics analysis

For quantitative profiling of metabolites in whole cell lysates and cell culture media, aliquots were prepared for polar metabolites, acylcarnitine profiling, and protein quantification, then stored at −80°C. Sulfur-containing metabolites, creatinine, S-adenosylmethionine, and S-adenosylhomocysteine were quantified using previously described methods [40, 41]. Lactate, TCA and glycolysis intermediates, organic acids, folates, amino acids, and neurotransmitters were profiled following established protocols [42]. Amino acid calibration curves were generated from a standardized mixture, while other metabolites were calibrated from in-house stock solutions. Accuracy was confirmed by monitoring specific metabolites in external quality controls and validated plasma samples, with data collected on a Sciex 6500+ ESI-tripleQ MS/MS coupled to a Nexera ultra-performance liquid chromatograph. Acylcarnitines were extracted from tissue homogenate and categorized by chain length, analyzed via a Sciex 5500+ MS/MS. Quantification was performed with Analyst® 1.7.2 software.

### Tumor interstitial fluid preparation for metabolomics

For the isolation of tumor interstitial fluids, tumors were excised from the animals as quickly as possible, washed with PBS, and then completely blotted dry. The tumors were subsequently centrifuged at 200g for 10 minutes at 4°C using an EASYstrainer™ (20 µm, Greiner BIO-ONE). Isolated interstitial fluid was stored at −80°C, and sulfur-containing metabolites were quantified as previously described [40, 41]. Lactate, TCA and glycolysis intermediates and other organic acids, folates and amino acids were determined as described in previous work [42, 43]. Folates, TCA intermediates, and amino acids (including urea cycle intermediates) were calibrated with a standardized amino acid mixture. For extraction, 20 µL fluid was treated with DTT, incubated, and processed with 0.1% formic acid in methanol. Calibration was validated with external quality controls, and quantification was carried out using Analyst® 1.7.2 software on an LC-MS/MS system.

### Single-cell sequencing

Alignment and quantification of single-cell RNA-seq data were performed using CeleScope (Singleron Biotechnologies, version 1.14.0) and aligned to the mouse genome “Mus_musculus_ensembl_92” (Ensembl). Count matrices were analyzed with Scanpy (version 1.9.1) [44]. Quality control excluded cells with fewer than 500 genes, more than 25,000 UMI counts, or over 10% mitochondrial counts. Gene expression was log-normalized and scaled by 10,000 [45]. PCA was performed on the top 5,000 variable genes, followed by Harmony integration to correct batch effects [46]. Leiden clustering (resolution 0.5) was used, and UMAP was employed for visualization [47, 48]. Differentially expressed genes (DEGs) were identified using Wilcoxon rank-sum test with adjusted p-values <= 0.05 and log fold change >= 1. GO enrichment analysis was performed with ClusterProfiler in R [37, 49, 50] (version 4.2.2) and GSEAPY for curated gene sets [51] in scRNA-seq data, visualized in UMAP.

### Embedding and preparation of pancreatic tissue for electron microscopy

On Day 1, pancreatic tissue was fixed in 4% paraformaldehyde and 2.5% glutaraldehyde, followed by washes and osmication. After dehydration, the tissue was incubated in uranyl acetate overnight. On Day 2, the tissue underwent ethanol treatments, propylene oxide immersion, and embedding in Durcupan. On Day 3, the tissue was cured, sectioned, and stained. Semi-thin (300 μm) sections were stained with toluidine blue, while ultra-thin (58 nm) sections were contrasted with lead citrate. Samples were sectioned on a Leica Ultracut UC6 and analyzed with a Zeiss LEO 906E transmission electron microscope

### Statistical analysis

Statistical analysis was conducted using GraphPad PRISM 9.2.0 software. For experiments analyzing only one variable across multiple conditions, ordinary one-way ANOVA multiple comparisons were employed. Groups or samples were considered significantly different when P < 0.05. Detailed statistical information for each experiment is provided in the corresponding supplementary Information. Large dimensional data from omics experiments were examined with R Studio using the indicated R packages [52].

### Grammatical and Stylistic Editing

To ensure clarity and precision, grammatical and stylistic revisions were made with the assistance of an AI-based language model (ChatGPT, OpenAI), which was used exclusively for language-related support. No content-related changes or data analysis were performed using this tool.

## Results

### In vitro screening reveals mitochondrial respiration as a common dependency in response to dual SHP2/MEK inhibition

We selected a panel of PDAC cell lines to screen for metabolism associated subtype-specific vulnerabilities upon SHP2/MEK inhibition *in vitro*. The human cell lines MIA PaCa II [*KRAS^G12C^, TP53^R248W^*], PANC-1 [*KRAS^G12D^, TP53^R273H^*] and YAPC [*KRAS^G12V^, TP53^H179R^*] were utilized, as well as a number of murine cell lines established from endogenous genetic tumors of the autochthonous *Kras^G12D/+;^ Trp53^fl/fl^; Ptf1a-Cre* (KPC) PDAC model. Cell lines were classified and chosen according to their molecular subtype based on morphology, protein expression profile and transcriptomic gene set enrichment, to guarantee for representative, cross-species coverage of the basal-to-classical PDAC subtype spectrum (Fig. 1*A* and Supplementary Fig. 1A-E).

**Figure 1:**
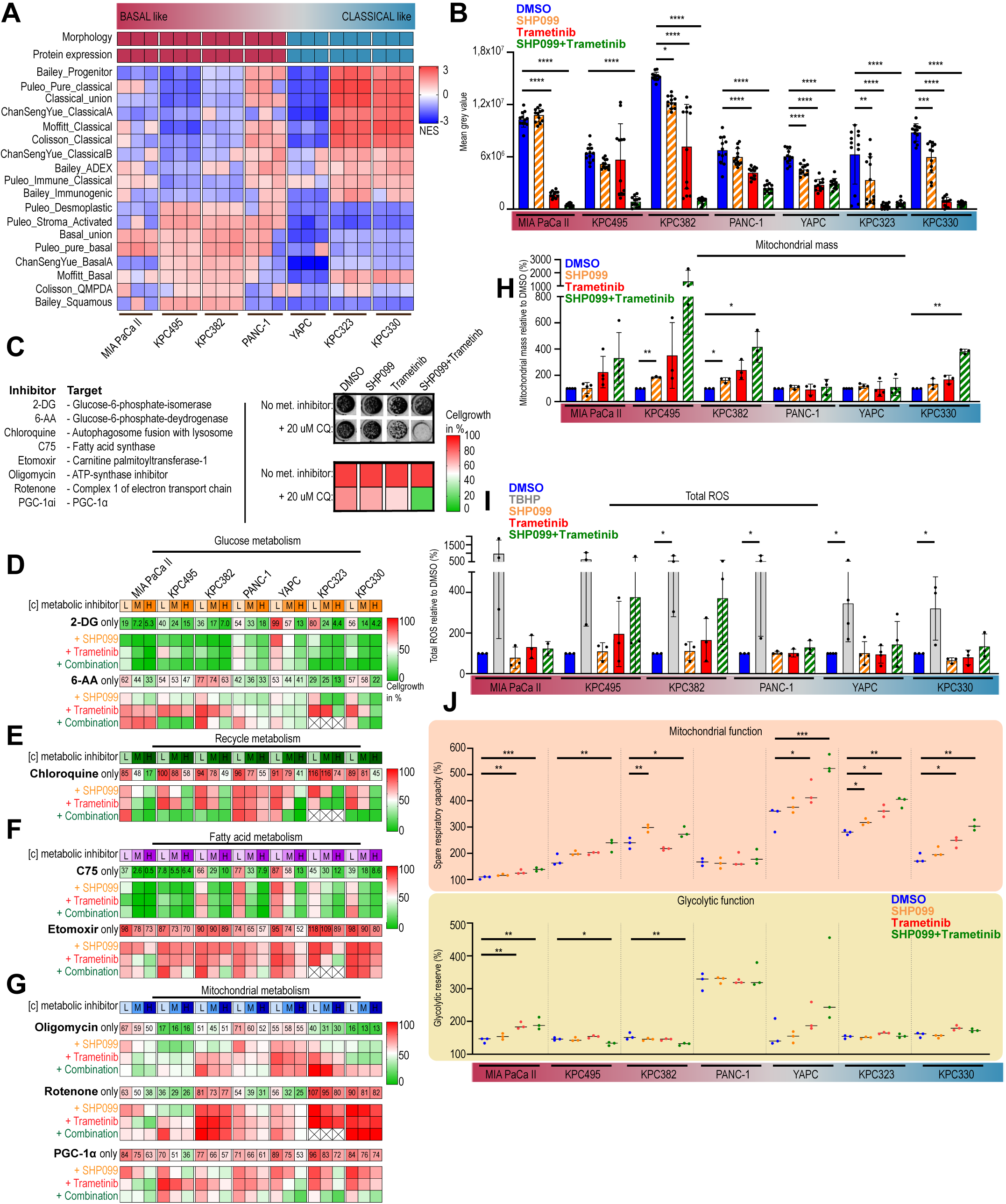
Pharmacological SHP2 and/or MEK inhibition impairs tumor cell metabolism. **(*A*)** Transcriptomic classification of human and murine PDAC cells into basal-like or classical-like subtypes. (***B*)** Cell proliferation of PDAC cells treated with DMSO, SHP099, Trametinib, or both. Data represent SD from 12 wells in one experiment. **(*C*) Left:** Metabolic inhibitors combined with MAPK inhibition. **Right:** Example proliferation assay, with color-coded MAPK treatment comparisons at ∼90% confluency with or without metabolic inhibitors. **(*D-G*)** Relative cell proliferation under MAPK inhibition plus increasing metabolic inhibitor doses in PDAC cells. Heatmap represents mean of 8 wells across two independent experiments; concentrations are depicted in Supplementary Fig. 2. **(*H*)** Mitochondrial mass by flow cytometry in PDAC cells, mean of 3–4 independent experiments. **(I**) Intracellular ROS by flow cytometry, with tert-Butyl hydroperoxide (TBHP) as ROS-inducing control; mean of 2–4 independent experiments. **(*J*)** Spare respiratory capacity and glycolytic reserve in PDAC cells, assessed via ECAR and OCR in 3 experiments with 6 technical replicates each. Statistical significance was determined via one-way ANOVA in panels B, H, I, J, with comparisons made against corresponding DMSO controls.* P < 0.05, ** P < 0.01, *** P < 0.001, **** P < 0.0001

The allosteric SHP2 inhibitor SHP099 demonstrated only minimal effect on proliferation, while the MEK1/2 inhibitor Trametinib separated cell lines into primary sensitive (Mia PaCa II, KPC323, KPC330) and intrinsic/rapid-adaptive resistant ones (KPC382, KPC495, PANC-1, YAPC) (Fig. 1*B*). Dual SHP099/Trametinib therapy enhanced sensitivity or overcame intrinsic resistance, with no subtype-specific response observed, suggesting broad applicability and therapeutic value for dual SHP2/MEK inhibition (Fig. 1*B*).

To gain insight into potential dependencies on specific metabolic pathways, cell lines were then exposed to various metabolic inhibitors with or without MAPK inhibition. (Fig. 1*C*, left). After allowing each MAPK treatment (without metabolic inhibitor) to reach full growth, we compared it to the same treatment with the metabolic inhibitor. This normalization enabled us to isolate the specific impact of metabolic inhibition on top of each MAPK therapy (exemplified in Fig. 1*C*, right).

Glycolysis inhibition using 2-Deoxy-D-glucose (2-DG) indicated higher sensitivity for basal-like cell lines. Adding SHP2 and/or MEK inhibition further sensitized classical murine cell lines without altering the high sensitivity of basal-like lines (Fig. 1*D*, top panel; supplementary Fig. 2).

Pentose phosphate pathway (PPP) inhibition via 6-Aminonicotinamide (6-AA) revealed heterogeneous results, with either increased or decreased PPP dependency across subtypes (Fig. 1*D*, bottom panel; supplementary Fig. 2).

Dependency of PDAC on autophagy, particularly after MAPK inhibition, was confirmed in 6 out of 7 cell lines using Chloroquine (CQ). Dual SHP2/MEK inhibition further potentiated this effect, increasing CQ sensitivity compared to Trametinib alone, irrespective of molecular subtype (Fig. 1*E*, supplementary Fig. 2). Fatty acid synthase (FAS) inhibition with C75 impaired proliferation across all cell lines but did not interact with SHP2 or MEK inhibition (Fig. 1*F*, supplementary Fig. 2). However, fatty acid oxidation (FAO) interference via Etomoxir increased sensitivity predominantly in rather basal-like cell lines (MIA PaCa II, KPC495, KPC382, and PANC-1), although off-target mitochondrial effects from Etomoxir concentrations [53] might have influenced the results.

Further screening indicated mitochondrial respiration changes, particularly with dual SHP2/MEK inhibition (Fig. 1*G*, top panel, Supplementary Fig. 2). Trametinib dampened ATP-synthase inhibitor oligomycin-induced proliferation impairment in 2/3 classical and 2/3 basal-like lines, with a more pronounced effect under combination therapy. Similar patterns were obtained with Rotenone, a complex 1 inhibitor, where Trametinib reduced its potency in 3/3 basal-like and 1/3 classical lines, further enhanced with dual inhibition (Fig. 1*G*, middle panel, Supplementary Fig. 2). PGC-1α inhibition showed increased sensitivity under dual inhibition across subtypes, further supporting a central role of mitochondrial metabolism (Fig. 1*G*, bottom panel, Supplementary Fig. 2).

Given the screening data pointing to mitochondria as a dominant metabolic target in response to dual SHP2/MEK inhibition across PDAC subtypes, we investigated mitochondrial behavior under RAS-pathway inhibition in more detail. Dual SHP099/Trametinib therapy increased mitochondrial mass in basal and classical lines, while reactive oxygen species (ROS) levels trended upward, albeit not significantly (Fig. 1*H* and 1*I*). Electron microscopy indicated changes in mitochondrial number and microstructure, particularly in the combination therapy, yet without a fully consistent pattern (supplementary Fig.3). Then, in order to gain insight into mitochondrial respiratory function, oxygen and proton flux analysis by Seahorse assays was performed. Combination therapy significantly increased spare respiratory capacity, describing the ratio of basal to maximum respiration and indicating enhanced mitochondrial fitness, in most cell lines (Fig. 1*J*, top panel). Basal respiration decreased in 2/3 basal-like lines (Supplementary Fig. 4A, top), while maximal respiration increased notably in classical-like lines (Supplementary Fig. 4A, middle), likely driving enhanced respiratory capacity in the classical-like group. Proton leak analysis showed no notable differences in oxidative phosphorylation (OXPHOS) efficiency between cell lines or treatments (Supplementary Fig. 4A, bottom). As mutated *KRAS* influences glycolysis, we assessed this pathway under MAPK inhibition. Seahorse assays revealed no significant changes in glycolytic reserve (Fig. 1*J*, bottom) or basal glycolysis between basal-like and classical cells (Supplementary Fig. 4B).

Altogether, these *in vitro* data suggest that indirect interference with RAS-activity via dual SHP2/MEK inhibition leads to substantial metabolic changes in PDAC tumor cells. While increased dependence on autophagy is further amplified in comparison to MEK1/2 inhibition alone, metabolic alterations are particularly linked to changes in mitochondrial abundance and function, across molecular subtypes.

### Multiomics confirm mitochondrial alterations in response to SHP2/MEK inhibition

To investigate the underlying properties of the screening response of our cell lines, we performed metabolomic, proteomic, and transcriptomic analyses. The human cell lines MIA PaCa II, PANC-1, and YAPC underwent mass spectrometric analysis of intra-/extracellular metabolites and targeted proteomics.

Metabolite analysis of the extracellular compartment revealed substantial alterations following therapy (Fig. 2*A*, supplementary Fig. 5*A-C*), particularly after 72 hours and most prominently with SHP2/MEK inhibition. Amino acids were the primary contributors to distinct metabolite profiles across treatments (Fig. 2*B*, supplementary Fig. 5*D-F*), with combination therapy significantly increasing extracellular amino acid levels (Fig. 2*C*, data for other cell lines not shown). Time-course analysis suggested these elevated levels stemmed from reduced uptake rather than increased secretion, as shown by serine and valine in MIA PaCa II cells (Fig. 2*D*). An increased abundance of amino thiols and neurotransmitters was also observed in the extracellular milieu (Fig. 2*B*, supplementary Fig. 5*D-F*). Given the pronounced mitochondrial alterations under combination therapy, we focused on metabolites linked to mitochondrial and ROS metabolism. In MIA PaCa II cells, glutathione, S-adenosylhomocysteine, cystathionine, methionine sulfoxide, malate, and citrate decreased intracellularly but increased extracellularly with combination therapy (Fig. 2*E*). Similar trends were seen in PANC-1 and YAPC cells, though less pronounced. Lipid metabolism also shifted, with an increase in acyl carnitines in MIA PaCa II cells across all treatments, including SHP099 monotherapy (Fig. 2*F*).

**Figure 2:**
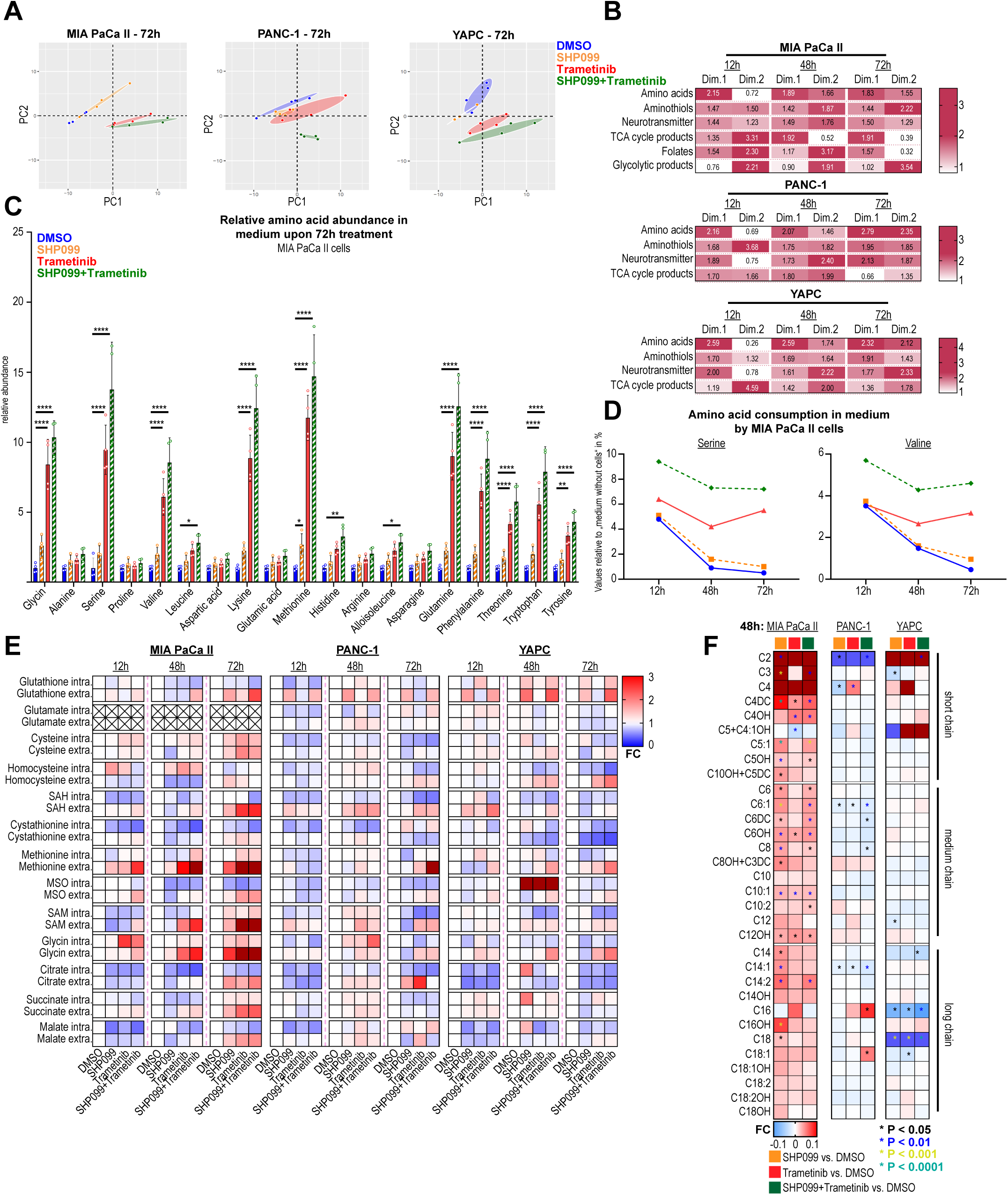
Pharmacological SHP2 and/or MEK inhibition influences intra- and extracellular metabolic composition. **(*A*)** PCA of extracellular metabolites in PDAC cell lines treated for 72 hours with DMSO, SHP099, Trametinib, or the combination (4 samples per treatment; 62 metabolites for MIA PaCa II, 54 for PANC-1 and YAPC). **(B)** Summarized PCA contributions factors that indicate how much each variable influences the principal components. PCA contribution of human PDAC cell lines at 12, 48, and 72 hours of treatment, showing variable loadings on the first two principal components, which capture the most variance. Values for the various metabolic pathways (e.g., amino acids) represent the mean of individual metabolites shown in Supplementary Figure 5. **(*C*)** Extracellular amino acid abundance in MIA PaCa II cells after 72 hours of treatment. **(*D*)** Three representative extracellular amino acids in MIA PaCa II relative to medium control. **(*E*)** Heatmap of intra- and extracellular metabolite levels in cells treated with SHP099, Trametinib, or both vs. DMSO controls (set to 1). Color gradient shows fold changes, with dark red indicating values >3 for clarity. **(*F*)** Acylcarnitine levels in PDAC cell lines after 48-hour treatments. Values >0.1 in dark red, <−0.1 in dark blue for clarity. Statistical significance was determined via one-way ANOVA in panel C, with comparisons made against corresponding DMSO controls. * P < 0.05, ** P < 0.01, *** P < 0.001, **** P < 0.0001. For panel F statistical analysis was performed with R [36] and the R package limma [52]. * P < 0.05, ** P < 0.01, *** P < 0.001, **** P < 0.0001.

Targeted proteomics revealed minimal effects of SHP099 monotherapy (supplementary Fig. 6*A-C*), while Trametinib had a stronger impact, most pronounced with combination therapy. Proteins involved in mitochondrial metabolism increased in MIA-PaCa II and PANC-1 cells following combination therapy (supplementary Fig. 6*D*), with mitochondrial fusion/fission proteins upregulated. Ferroptosis regulators also showed changes: GPX4 expression was downregulated in PANC-1 and YAPC under Trametinib and combination therapy. Other ferroptosis-related proteins exhibited notable changes, such as decreased ferroptosis suppressor protein 1 in MIA-PaCa II and YAPC, and increased long-chain-fatty-acid-CoA ligase 4 and transferrin receptor protein 1 in YAPC. Collectively, metabolomics and proteomics confirmed dual SHP2/MEK inhibition affects proliferation-related pathways, mitochondrial dynamics, and ferroptosis regulation.

For RNA-sequencing, we integrated KPC mouse cell lines, expanding the screening collection with additional lines to ensure dense representation of murine PDAC subtypes (Supplementary Fig. 1A, C, E). SHP099 monotherapy minimally impacted gene expression (Fig. 3*A*), while Trametinib and combination therapy showed greater influence. Gene set enrichment analysis (GSEA) highlighted metabolic gene alterations (Fig. 3*B*), revealing downregulation of DNA/RNA and amino acid metabolism genes. Combination therapy uniquely upregulated lipid and steroid metabolism-related genes. Downregulation of glycolysis-related gene sets under Trametinib and combination therapy confirmed KRAS dependency, while SHP2 inhibition had little effect. Autophagy-related gene sets were upregulated with combination therapy in nearly all cell lines, compared to fewer under Trametinib alone, suggesting sustained MAPK inhibition (Fig. 1*E*). Significant changes in OXPHOS and ROS-associated pathways were also observed, particularly under combination therapy. ROS protective gene sets, including glutathione and cysteine/methionine metabolism, were downregulated in most murine cell lines with combination therapy (combined comparisons with DMSO and Trametinib).

**Figure 3:**
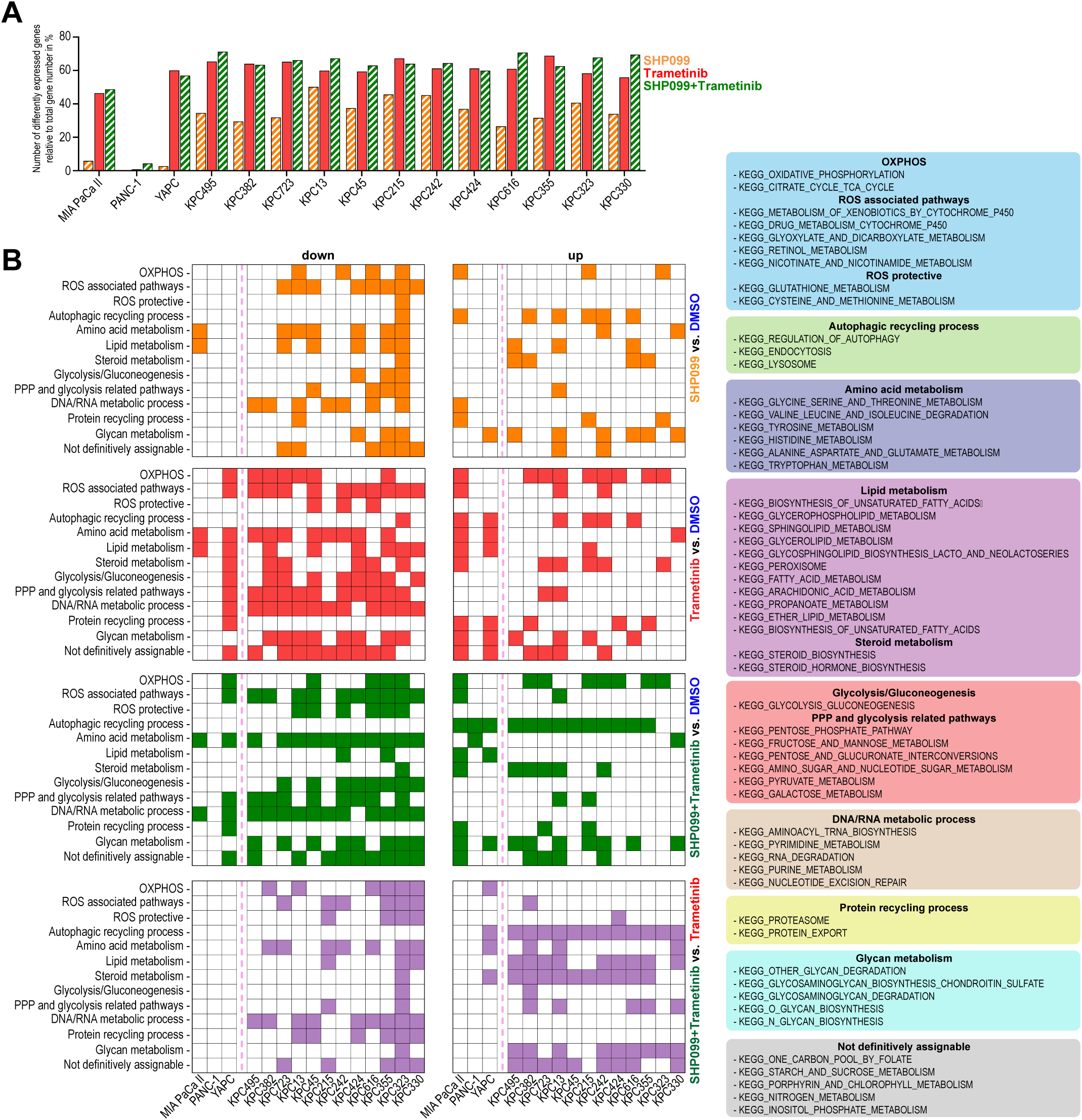
Transcriptomic analysis reveals metabolic adaptations in tumor cells with SHP2 and/or MEK inhibition. **(*A*)** Number of differentially expressed genes (DEGs) in human PDAC and murine KPC cells treated with SHP099, Trametinib, or both; DEGs with adj. P < 0.01 are shown. **(*B*)** KEGG-based gene set enrichment analysis (GSEA) of human and murine cells treated for 48 hours with DMSO, SHP099, Trametinib, or their combination. Each colored square represents a significant upregulation or downregulation of at least one of gene set, as depicted from the right. "Not definitively assignable" indicating unclear metabolic classification. Gene sets with adj. P < 0.25 are indicated.

In sum, RNA-sequencing analysis corroborated our functional observations regarding anabolism, glycolysis, autophagy, and mitochondrial function, without a clear correlation with molecular PDAC subtypes.

### Genetic knockout of SHP2 has a profound impact but does not recapitulate or inform about pharmacologic SHP2 inhibition

With the aim to dissect the particular contribution of SHP2 to PDAC metabolism in more detail, we generated *PTPN11* knockout (KO) cell lines. For human cell lines CRISPR-Cas9 technology was employed, as murine counterparts we used cell lines derived from our previously reported dual recombinase PDAC mouse model [22].

Knockout was confirmed by Western Blot. Analysis of core signaling demonstrated increased KRAS levels, higher basal pERK levels and/or pAKT and pSTAT3 levels in CRISPR-KO cells (supplementary Fig. 7*A*) as a sign of compensation for loss of SHP2. Yet, proliferation was impaired more dramatically in *PTPN11* KOs as compared to SHP099 treatment (supplementary Fig. 7*B*).

The KOs also exhibited a very different metabolic behavior compared to parental cells in response to pharmacological SHP2 inhibition. Specifically, MIA-PaCa II KO cells displayed heightened sensitivity to OXPHOS inhibition relative to untreated or SHP099-treated parentals (supplementary Fig. 7*C*). Additionally, these cells exhibited an increased mitochondrial mass, respiratory capacity but decreased ROS levels (supplementary Fig. 7*D*, *E* & *G*). Notably, KO cell lines were susceptible to FAO inhibition by Etomoxir (supplementary Fig. 7*C*), which resembled more closely the behavior of cell lines treated with combination therapy as shown in Figure 1*F*.

Measurement of extracellular metabolites revealed a distinct metabolic shift separating KO cell lines from SHP2-proficient cells and pharmacological inhibition (Supplementary Fig. 10F). Transcriptional analysis showed a shift in subtype classification after PTPN11 KO (Supplementary Fig. 7H). The murine KO cell lines F2453 and F2683 were largely similar, with F2683 showing the best functional compensation for SHP2 loss (Supplementary Fig. 7I-K).

In summary, genetic SHP2 knockout induces more profound metabolic and signaling changes than pharmacological inhibition, likely due to clonal selection during the KO process. Thus, PTPN11 KO cell lines may not directly reflect pharmacologic SHP2 inhibition outcomes.

### Mitochondrial adaptions are sustained in therapy-resistant cell lines

We investigated therapy-resistant states by treating cell lines with SHP099, Trametinib, or their combination for at least 8 weeks, leading to adaptive resistance. KRAS signaling was upregulated in MIA PaCa II and PANC-1 cells, along with E2F target genes and IL-6-JAK-STAT3 response (Supplementary Fig. 8A). In YAPC, KRAS upregulation and early activation of compensatory pathways like IL2-STAT5 and IL6-JAK-STAT3 were observed already after short-term treatment (Supplementary Fig. 8A).

The increased mitochondrial mass observed during short-term combination therapy (Fig. 1*H*) also became significantly evident in 2/3 of basal cell lines during long-term therapy (Supplementary Fig. 8B), while classical lines tended to show reduced mitochondrial mass (Supplementary Fig. 8B). Intracellular ROS levels were notably elevated in treated MIA PaCa II, while no significant changes were observed in the other cell lines (Supplementary Fig. 8C).

The resistant cell lines displayed unchanged sensitivity to glycolysis inhibition but increased dependency on the PPP, with 6/7 lines showing strong sensitivity to 6-AA under combination therapy, unlike short term treatments (Supplementary Fig. 8D and Fig. 1*D*). Sensitivity to autophagy inhibition by chloroquine persisted, particularly in combination therapy, affecting 5/7 cell lines (Supplementary Fig. 8E and Fig. 1*E*). FAS inhibition sensitivity by C75 remained unchanged, while the effect of Etomoxir was enhanced in 2/7 Trametinib and combination resistant lines. All resistant lines (7/7) showed increased sensitivity to Rotenone, and 4/7 to Oligomycin (Supplementary Fig. 8G).

Interestingly, long-term treatment with Trametinib or the combination of SHP099 and Trametinib resulted in downregulation of OXPHOS-related gene transcription in all three human cell lines (Supplementary Fig. 8H). Additionally, MIA PaCa II cells showed downregulation of the citric acid cycle and 2/3 of human cell lines exhibited downregulation of the ROS-protective glutathione metabolism.

Together, these data from long-term treated cell lines suggest a persisting impact of SHP2/MEK inhibition on mitochondrial metabolism even in a therapy-resistant state.

### Mitochondrial adaption following dual SHP2/MEK inhibition is confirmed *in vivo*

To see if the observed metabolic properties following SHP2/MEK inhibition prove true *in vivo*, we treated KPC mice, monitored tumor volumes using magnetic resonance imaging (MRI) and collected tumor interstitial fluid (TIF). The tumors used for TIF isolation displayed the previously described [22] pattern of response to SHP2/MEK inhibition in terms of tumor dynamics (Fig. 4*A*).

**Figure 4:**
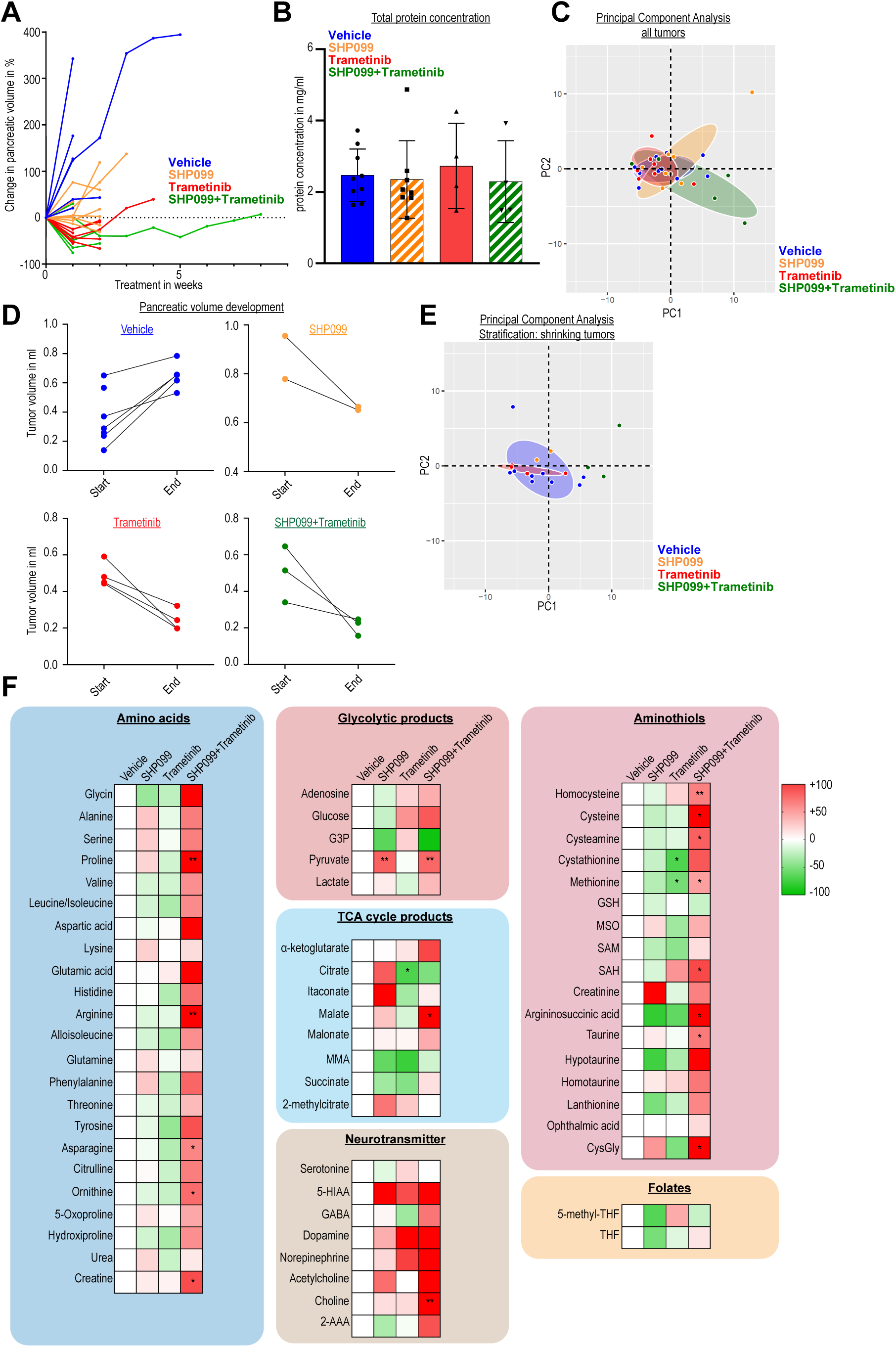
Tumor interstitial fluids reveal metabolic adaptations in amino acid metabolism and ROS-associated pathways with dual SHP2 and MEK inhibition in KPC tumors. **(*A*)** Dynamics of KPC tumors in mice treated with vehicle (control), SHP099 (75 mg/kg), Trametinib (1 mg/kg), or their combination. Tumor interstitial fluids (TIFs) were analyzed via LC-MS/MS. **(*B*)** TIF protein concentration in different therapy groups. **(*C*)** PCA of 63 metabolites from TIFs in KPC tumors under therapy. **(*D*)** PDAC tumor development of shrinking tumors using magnetic resonance imaging. **(*E*)** PCA of tumors stratified by shrinkage behavior. **(*F*)** Metabolite abundance under targeted therapy relative to vehicle-treated tumors, organized by metabolic pathways. Supplementary Fig. 9 details individual metabolites. Statistical significance was determined using one-way ANOVA in panel F, with comparisons made against corresponding DMSO controls* P < 0.05, ** P < 0.01. Abbreviations: G3P – glyceraldehyde-3-phosphate, MMA – methylmalonic acid, 5-HIAA – 5-hydroxyindoleacetic acid, 2-AAA – 2-aminoadipic acid, GSH – glutathione, MSO – methionine sulfoximine, SAM – S-adenosylmethionine, SAH – S-adenosylhomocysteine, CysGly – cysteinylglycine, THF – tetrahydrofolate

Metabolite profiling of TIFs by LC-MS/MS revealed no significant differences in protein concentration (Fig. 4*B*). While PCA of the full sample set did not show clear clustering by treatment, three combination-treated tumors formed a distinct subcluster, correlating with tumor volume reduction (Fig. 4*C*, *D*). The other two combination samples were either treated for a prolonged period (55 days) or did not respond to the treatment in the same manner.

To maintain consistency, we selected TIFs from tumors exhibiting volume regression during two weeks of therapy for further analysis (Fig. 4*D*). This refined PCA revealed separate clustering for combination-treated TIFs (Fig. 4*E*). Metabolite analysis showed elevated amino acid levels in combination-treated tumors, consistent with reduced uptake due to proliferation inhibition (Fig. 4*F*, supplementary Fig. 9). Glucose and glyceraldehyde 3-phosphate levels suggested reduced glycolysis in both Trametinib and combination therapy groups (Fig. 4*F*, supplementary Fig. 9). TCA cycle intermediates like alpha-ketoglutarate and citrate behaved similarly between treatments, although malate was significantly increased in the interstitial space of combination therapy samples (Fig. 4*F*, supplementary Fig. 9).

Interestingly, an increased abundance of neurotransmitters was found, particularly in TIFs from SHP099/Trametinib treated tumors (Fig. 4*F*, supplementary Fig. 9).

Finally, the composition of the aminothiols in TIFs indicated changes in mitochondrial respiration and ROS defense. Comparison of SHP099 and of Trametinib with combination treatment displayed striking differences, for instance, combination treatment resulted in significantly elevated levels of cysteine, methionine, S-adenosylmethionine (SAM) and SAH (Fig. 4*F*, supplementary Fig. 9). The upregulation of the methionine cycle and transsulfuration pathway may be elicited to counter the anti-proliferative and pro-oxidative effects of the combined treatment, respectively.

In summary, KPC tumors treated with dual SHP099/Trametinib therapy showed distinct metabolic changes in the interstitial space, particularly in mitochondrial respiration and ROS protection, compared to Trametinib alone.

To further substantiate our observations, we treated KPC mice and collected fresh samples after short-term and survival endpoint treatments, confirming tumor dynamics (Fig. 5*A*). Electron microscopy showed significantly larger mitochondria in tumors treated with SHP099/Trametinib compared to controls (Fig. 5*B*, left). Both short-term treatments increased mitochondrial size, but only SHP099/Trametinib had a persistent effect (Fig. 5*B*, right). Additionally, mitochondrial cristae were more densely packed, with intra- and intertumoral heterogeneity (Fig. 5*C*). These results aligned with our *in vitro* observations, reinforcing the conclusion that dual SHP2 and MEK inhibition induces significant mitochondrial alterations in PDAC, both *in vitro* and *in vivo*.

**Figure 5:**
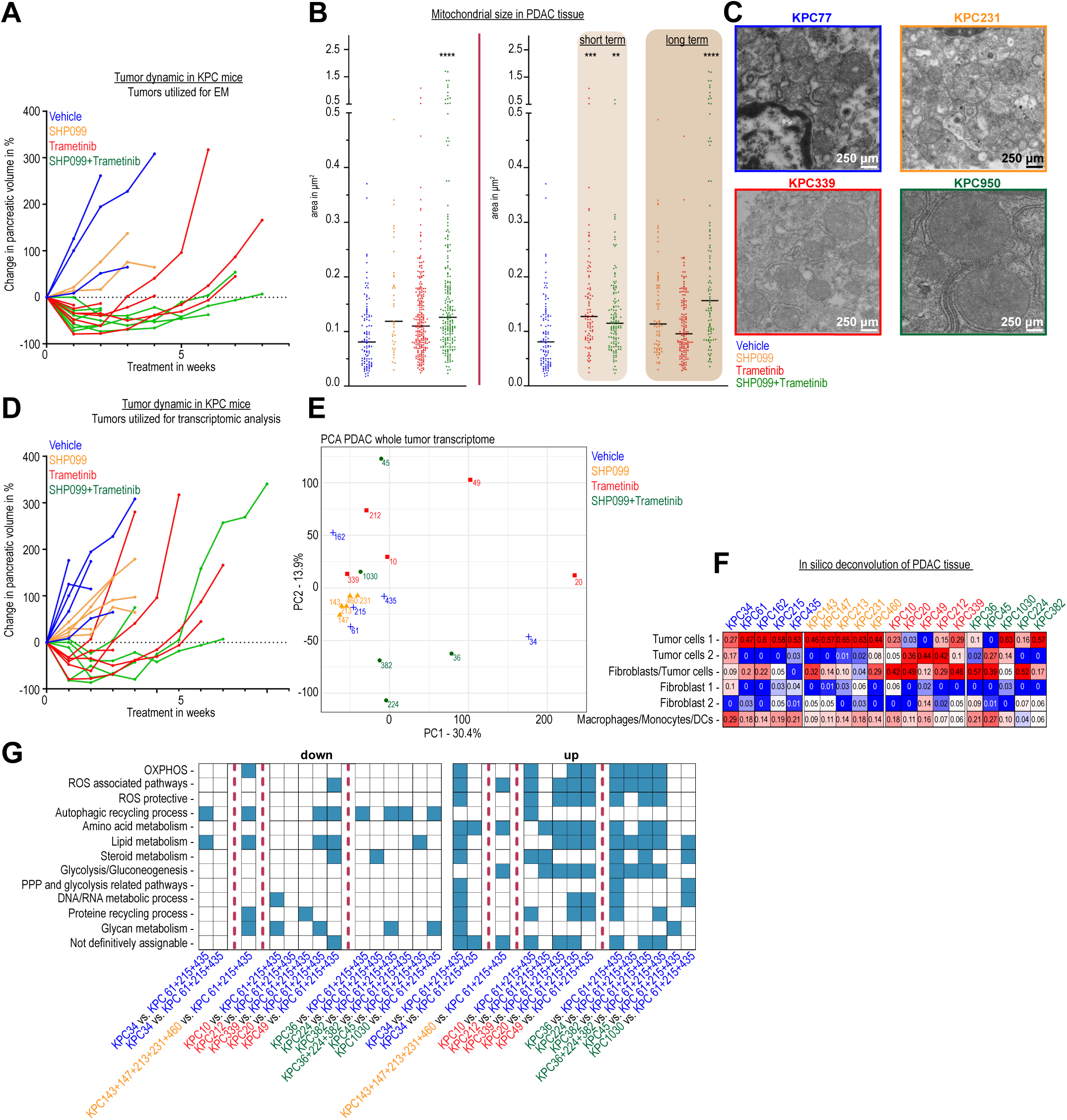
Dual SHP2 and MEK inhibition reveals mitochondrial adaptations in vivo. **(*A*)** Tumor dynamics in KPC mice treated with vehicle (control), SHP099 (75 mg/kg), Trametinib (1 mg/kg), or the combination, with tumors processed for electron microscopy. **(*B*) Left** Mitochondrial diameter in PDAC tumor cells from KPC mice, shown for all therapies. **Right** Separated by short (≤ 2 weeks) and long (> 2 weeks) therapy durations. **(*C*)** Representative mitochondrial morphology in KPC PDAC tumor cells under targeted therapy. **(*D*)** Tumor dynamics in KPC mice treated with vehicle (control), SHP099 (75 mg/kg), Trametinib (1 mg/kg), or the combination, with tumors processed for whole tissue RNA sequencing. **(*E*)** PCA of KPC tumors following therapy. **(*F*)** In silico deconvolution of sequenced KPC tumors. **(*G*)** KEGG-based GSEA of KPC tumors, showing significant gene set upregulation or downregulation, with gene sets of adj. P < 0.25 indicated. Statistical significance in panel B was determined by one-way ANOVA against vehicle controls: * P < 0.05, ** P < 0.01, *** P < 0.001, **** P < 0.0001.

We then performed bulk RNA-sequencing of tumor tissue from animals treated until survival endpoint (Fig. 5*D*). Principal component analysis (PCA) revealed distinct transcriptomic profiles, demonstrating inter-tumor heterogeneity, with three vehicle-treated samples (KPC61, KPC215, KPC435) clustering together as a reference for comparison (Fig. 5*E*). *In silico* deconvolution indicated that epithelial cells dominated bulk-Seq transcripts, followed by fibroblasts and myeloid cells (Fig. 5*F*), supporting conclusions mainly regarding tumor cell transcription. GSEA highlighted upregulation of OXPHOS-related pathways in Trametinib-treated (3/5) and combination-treated (4/6) samples, along with ROS-associated pathways (Fig. 5*G*).

To get more precise insight into cell specific transcriptional alterations in treated KPC tumors, we conducted single-cell RNA-sequencing of tumors from treated KPC mice. Tumor-cell clusters were identified and subclustered into groups with predominantly epithelial (E1-E5) and mesenchymal (M1-M3) features, corresponding to classical and basal-like transcriptional subtypes, respectively (Fig. 6*A*-*C*). Notably, mesenchymal clusters (M1-M3) were enriched in tumors after long-term Trametinib but less so after long-term dual SHP099/Trametinib (Fig. 6*D*).

**Figure 6:**
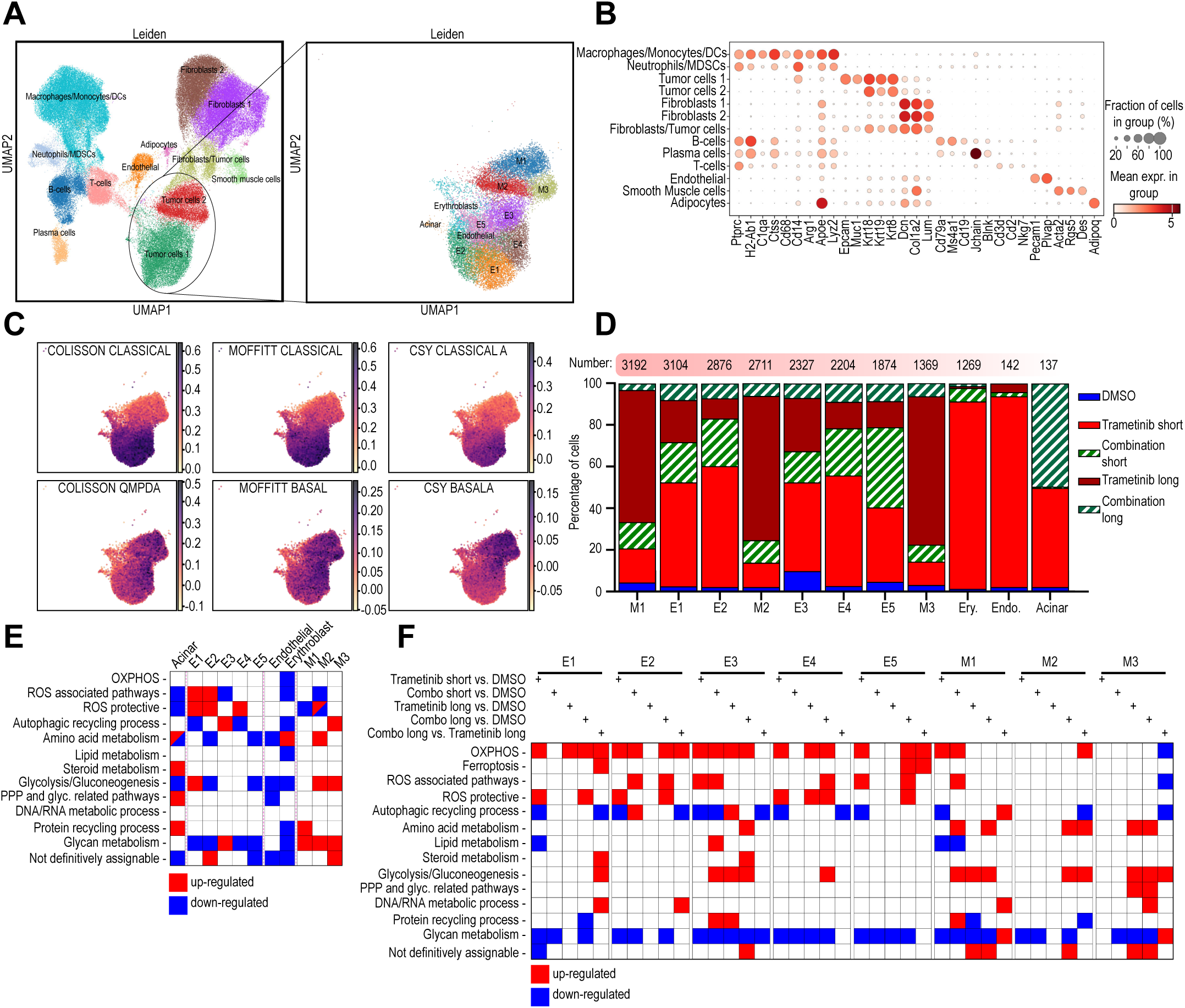
Single-cell transcriptomics reveals enhanced mitochondrial involvement following dual MAPK inhibition. (*A*) Left: Batch-corrected UMAP plot of the cells from PDAC tissue from individual KPC mice treated either with vehicle (acts as control), Trametinib (1mg/kg), or the combination of SHP099 (75mg/kg) and Trametinib. KPC mice treated ≤ 2 weeks (short) and > 2 weeks (long) were included. **Right:** UMAP plot highlighting the subclustering of the epithelial compartment. **(*B*)** Dot plot displaying the expression levels of selected genes across various cell types. **(*C*)** Subtype-specific contributions to the epithelial subclusters. **(*D*)** Percentage distribution of the indicated treatments across different epithelial subclusters. **(E)** Cluster-specific gene set enrichment analysis within the epithelial compartment, including the top 500 up- and downregulated differentially expressed genes. **(F)** Treatment-specific GSEA within the epithelial compartment, based on the top 500 up- and downregulated DEGs. Abbreviations: DCs – dendritic cells, MDSCs – myeloid-derived suppressor cells

Metabolic gene set analysis revealed dominant alterations in ROS associated pathways across the majority of tumor cell clusters (Fig. 6*E*). Combination therapy impacted transcription of OXPHOS gene sets in most tumor cell clusters, alongside ROS-associated and ROS-protective pathways (Fig. 6*F*).

Collectively, altered mitochondrial amount and function was confirmed *in vivo* in PDAC tumor cells, particularly following dual SHP2/MEK inhibition. The data also indicate a sustained and even more pronounced effect in terms of mitochondrial adaptations with long-term treatment conditions under which therapy resistant tumors have evolved.

### Patient-derived PDAC organoids confirm mitochondrial alterations in response to dual SHP2/MEK inhibition

To corroborate findings, we utilized patient-derived PDAC organoids (PDOs) from our own in-house reverse translational platform. Four PDOs with basal-like and four PDOs with classical subtype were selected (supplementary Fig. 10*A*).

While morphology of the selected organoid lines is pretty similar (supplementary Fig. 10*B*), transcriptional characteristics differ, with basal-like lines exhibiting enrichment for Epithelial-Mesenchymal Transition (EMT) and TGF-beta signaling, and downregulation of OXPHOS and fatty acid metabolism in a treatment naïve state (supplementary Fig. 10*C*). Interestingly, basal-like organoids appeared to be more sensitive to Trametinib and dual treatment (supplementary Fig. 10*D*).

The organoid lines most sensitive to glycolysis inhibition with 2-DG were of the basal-like subtype and for certain organoid lines (B150, B100) SHP2 and MEK inhibition markedly sensitized for glycolysis inhibition (supplementary Fig. 10*E*).

Unlike for 2D cell lines, combination therapy did not increase autophagy dependence of patient derived organoids (supplementary Fig. 10*E*).

However, Etomoxir was effective in virtually all classical and 50% of basal-like lines under SHP2/MEK inhibition (supplementary Fig. 10*E*).

Classical organoids showed higher sensitivity to PGC-1α inhibition upon dual therapy, with some in basal-like, as M7.

Inhibition of OXPHOS with Rotenone demonstrated increased sensitivity in four of the eight organoid lines and decreased sensitivity in one line (M5) in the presence of combination therapy (supplementary Fig. 10*E*). Similar, yet less striking effects were also observed in the combination therapy with Oligomycin (supplementary Fig. 10*E*). Response patterns did not correlate with transcriptional subtype.

Organoid lines display considerable variation across biological replicates. And, due to inherent culture limitations, readout of metabolic inhibitor screening (supplementary Fig. 10*E*) was performed at the same time for all conditions rather than allowing each to reach full confluency as in experiments with 2D cell lines. It is therefore possible that metabolic alterations were not fully captured.

Still, taken together, these data confirm mitochondrial alterations in response to dual SHP2/MEK inhibition in the majority of patient-derived PDAC organoid lines.

### An exploitable dependency on GPX4 evolves with dual SHP2/MEK inhibition

As *in vitro* analyses already suggested (Fig. 1*H* and 1*I*, 2*E* and supplementary Fig. 6*D*), mitochondrial alterations following dual SHP2/MEK inhibition appear to correlate with oxidative stress and lipid peroxidation within tumor cells.

To now measure membrane lipid peroxidation *in vivo* we performed malondialdehyde (MDA) immunohistochemistry on treated KPC tumor tissue slides. Histoscore quantification revealed significantly increased MDA staining levels in combination-treated tumors (Fig. 7*A*).

**Figure 7:**
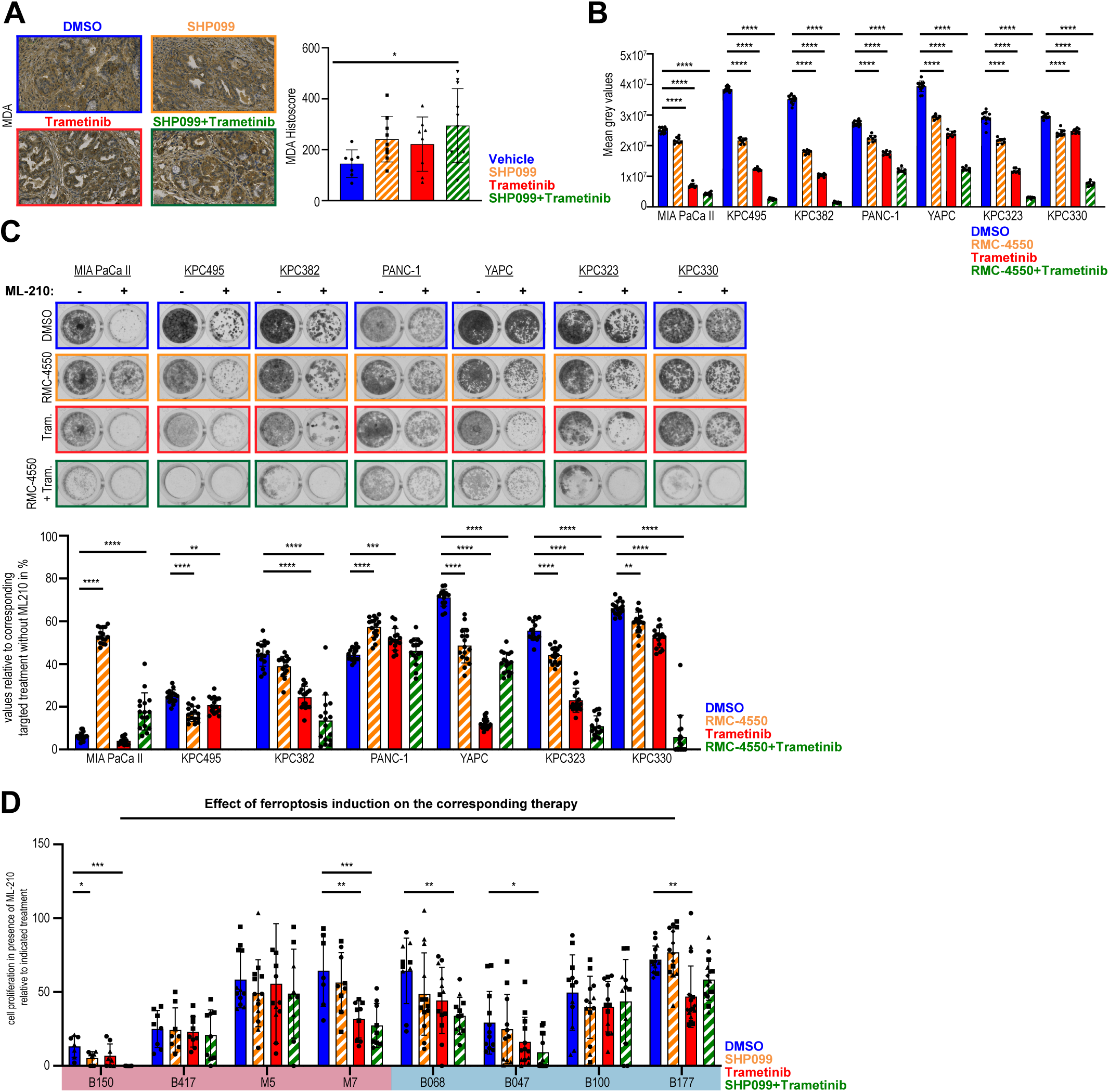
The combination therapy enhances sensitivity to ferroptosis induction. **(*A*) Left:** Malondialdehyde (MDA) staining; **Right**: MDA quantification in KPC mice treated with vehicle, SHP099 (75 mg/kg), Trametinib (1 mg/kg), or their combination. Data represent the mean of 5–10 tumor areas (comparable to those shown on the left) from histological sections. **(*B*)** Cell proliferation of human PDAC cell lines and murine KPC cells treated with DMSO, RMC-4550 (15 μM), Trametinib (10 nM), or both. Error bars show SD of 8 wells from one experiment. **(*C*)** Cell proliferation of human PDAC cell lines and murine KPC cells treated with DMSO, RMC-4550 (15 μM), Trametinib (10 nM), or both, plus GPX-4 inhibitor ML-210. Concentrations of ML-210 were 2 μM for MIA PaCa II, KPC495, and KPC382 cells; 10 μM for YAPC, KPC323, and KPC330 cells; 0.25 μM for PANC-1 cells. Error bars represent SD of 16 wells from one experiment. **(*D*)** Relative organoid proliferation with or without MAPK inhibition (DMSO, SHP099 (15 μM), Trametinib (25nM), or their combination) plus ML-210. The data illustrate the effect of ML-210 on top of each MAPK inhibition. Data normalized to MAPK inhibition alone (e.g., SHP099+ML-210 vs. SHP099). Data shown from 1-3 biological replicates (3-5 samples each). Statistical significance (B, C, D) was assessed by one-way ANOVA against DMSO controls: * P < 0.05, ** P < 0.01, *** P < 0.001, **** P < 0.0001

These findings suggested enhanced vulnerability to ferroptosis induction via GPX4 inhibition. We therefore treated various cell lines with the SHP2 inhibitor RMC-4550, Trametinib, and the GPX4 inhibitor ML-210 (Fig. 7*B* and *C*). The response to the combination therapy with RMC-4550 was comparable to that observed with SHP099 (Fig. 7*B*). Induction of ferroptotic cell death by targeting GPX4 was facilitated in the context of combined SHP2/MEK inhibition in all cell lines (Fig. 7*C*).

Notably, MIA PaCa II, KPC495, KPC382, KPC323 and KPC 330 exhibited minimal, if any, remaining viable cells following the triple treatment (Fig. 7*C*).

Consistent with previous findings, dual SHP2/MEK inhibition profoundly increased sensitivity to GPX4 interference, as evidenced by the further reduced proliferation of the patient-derived organoids from supplementary figure 10D when GPX4 inhibition was added (Fig. 7D).

Together, these data suggest a therapeutically exploitable and subtype-independent vulnerability to induction of ferroptotic cell death that evolves in PDAC in response to dual SHP2/MEK inhibition.

## Discussion

Even considering the burgeoning field of direct RAS inhibition, pancreatic adenocarcinoma (PDAC) remains among the cancers the most difficult to treat. New orally bioavailable allosteric SHP2 inhibitors have demonstrated promising potential, and have entered clinical evaluation in vertical RAS-pathway inhibition combinations. While addition of allosteric SHP2 inhibitors to MEK, ERK or RAS-targeted drugs yields synergism and potently delays resistance development, therapy failure needs to be anticipated and inadvertently ensues even with these combination strategies.

Previous studies have described oncogenic KRAS-driven metabolic rewiring [9–11, 54–58] and metabolic effects of MEK inhibition in PDAC [59], but the precise metabolic implications of vertical RAS-pathway inhibition including SHP2 as a target have not been evaluated. We here aimed to identify metabolic vulnerabilities emerging with deep inhibition of the RAS-MAPK-signaling pathway employing dual SHP2/MEK inhibition in PDAC, building on our previous work [22].

We here use PDAC subtype spectrum representative cell lines, autochthonous in vivo models and patient derived organoids, highlighting multilayered metabolic alterations in response to dual SHP2/MEK inhibition.

Our data indicate that basal-like cell lines rely more on glycolysis compared to classical cell lines (Fig. 1*D*), aligning with literature linking the basal subtype to glycolytic metabolism [21]. Yet, an overall reduction in glycolysis and/or PPP activity with MEK inhibition and combination therapy was observed. All cell lines analyzed demonstrated sensitivity to glycolysis inhibition and corresponding transcriptomic alterations, highlighting a conserved dependency of PDAC on glycolytic pathways, consistent with KRAS signaling supporting anabolic glucose metabolism [10, 11, 55, 60].

PDAC cells typically upregulate autophagy in response to MEK1/2 or ERK1/2 inhibition [55, 59]. We here find that addition of allosteric SHP2 inhibition enhances this response, an effect remaining with long-term treatment. Transcriptomic data confirmed an upregulation of autophagy-related genes with combination therapy (Fig. 3*B*). Dual SHP2/MEK inhibition in combination with autophagy inhibition may therefore be a more promising strategy for clinical application. Notably, a previous study reported that allosteric SHP2 inhibitors could inhibit autophagy due to their accumulation in lysosomes [61].

Our results also reveal reduced DNA/RNA metabolic processes and decreased amino acid influx in treated tumor cells (Fig. 2, Fig. 3*B*, and Fig. 4*F*). These effects are closely linked to inhibited proliferation but may influence cellular state and the tumor microenvironment (TME) with cancer-associated fibroblasts and various immune cells being involved. For instance, PDAC tumor cell glutamine uptake and metabolism has been demonstrated to uphold redox balance [9, 62, 63]. In the TME, elevated glutamine levels can enhance T cell survival, while MDSCs require glutamine for their immunosuppressive functions, indicating the complexity of metabolic processes in cancer tissues. [64–66].

Crucially, our data show a pronounced impact of RAS-pathway inhibition on mitochondrial dynamics both *in vitro* and *in vivo*, persisting into therapy resistant states. Dual SHP2/MEK inhibition leads to notable changes in mitochondrial metabolism (Fig. 1*H*&*J*, 3*B*, 4*F*, 5*B*&*G*). Tumor cells alter mitochondrial activity in response to combination therapy, as indicated by increased mitochondrial mass and elevated ROS levels esp. in basal-like cell lines. Electron microscopy imaging and increased spare respiratory capacity support these findings. Analyses of *in vivo* TIFs reveal significant differences in aminothiol levels between Trametinib and combination therapy. Additionally, cysteine and methionine metabolism are downregulated in murine KPC cell lines (Fig. 3*B*). As these alterations bring about a dependency on lipid peroxidase pathways and an increased vulnerability for induction of ferroptotic cell death, targeting GPX4 proves efficient in reinforcing vertical RAS-pathway inhibition via SHP2/MEK. These observations are further supported by recent findings showing that ROS-associated pathways and ferroptosis protective mechanisms play an important role in the adaptive resistance continuum [67].

Our findings build on and connect two previously reported PDAC characteristics, 1^st^ RAS-signaling driving glutamine metabolism for ROS-homeostasis [9, 62, 63] and 2^nd^mitochondrial dependency upon genetic loss of the KRAS oncogene [55], with a more generally described therapy-resistant cancer cell state dependent on lipid peroxidase pathways [68, 69]. Further, our work expands the notion that PDAC homeostasis and growth requires down-regulation of ferroptosis [70].

We cannot exclude that parts of the effects observed also relate to dysregulated SHP2 activity specifically in mitochondria. Allosteric inhibition of SHP2 might disrupt phosphorylation patterns in the respiratory chain, possibly enhancing mitochondrial activity and ROS production following combination therapy [71–73].

In summary, our study demonstrates interference with RAS-signaling via combined SHP2/MEK inhibition provoking mitochondrial alterations *in vitro* and *in vivo*, irrespective of molecular PDAC subtype. This response is present already with short term dual treatment but also persists into a therapy resistant state evolving with long-term inhibition. These adaptations bring about a vulnerability to induction of ferroptotic cell death via lipid peroxidase inhibition, providing a metabolic lever to reinforce SHP2/MEK inhibition in particular, and likely RAS-directed targeted PDAC treatment in general.

## Grant support and assistance

This study was funded by the German Cancer Aid, Deutsche Krebshilfe, Project ID 70113697 (to D.A.R.) and by the German Research Foundation, Deutsche Forschungsgemeinschaft (DFG), CRC1479 (Project ID 441891347, P17 to D.A.R., P06 to O.G., S01 to M.B and O.S., S02 to W.R.). M.B. is supported by the DFG, within CRC1160 (Project ID 256073931, Z02), CRC/TRR167 (Project ID 259373024, Z01), CRC1453 (Project ID 431984000, S01), TRR 359 (Project ID 491676693, Z01), and FOR 5476 UcarE (Project ID 493802833, P07). M.B. also acknowledges funding from the German Federal Ministry of Education and Research (BMBF) within the Medical Informatics Funding Scheme, PM4Onco–FKZ 01ZZ2322A. G.A. is supported by the BMBF via EkoEstMed–FKZ 01ZZ2015. O.G. is supported by the DFG, within CRC1160 (Project ID 256073931, C02), CRC/TRR167 (Project ID 259373024, A08), CRC1425 (Project ID 422681845, P10), GRK2606 (Project ID 423813989), and by the European Research Council (ERC) through Starting Grant 337689, Proof-of-Concept Grant 966687, and the EU-H2020-MSCA-COFUND EURIdoc programme (No. 101034170). L.H. and O.G. are supported by the DFG under Germany’s Excellence Strategy (CIBSS – EXC-2189 – Project ID 390939984).

## Abbreviations

2-DG: 2-deoxy-D-glucose
6-AA: 6-aminonicotinamide
CQ: chloroquine
DEGs: differentially expressed genes
E: epithelial
EMT: epithelial-mesenchymal transition
ERK: extracellular signaling kinases
FAO: fatty acid oxidation
FAS: fatty acid synthase
GPX-4: glutathione peroxidase
KPC: KrasG12D/+; Trp53fl/fl; Ptf1a-Cre
KO: knockout
LC-MS/MS: liquid chromatography with tandem mass spectrometry
MDA: malondialdehyde
MRI: magnetic resonance imaging
M: mesenchymal
MSO: methionine sulfoxide
MAPK: mitogen-activated protein kinase
MEK: mitogen-activated protein kinase kinase
PDAC: pancreatic ductal adenocarcinoma
PDOs: patient-derived PDAC organoids
PPP: pentose phosphate pathway
PCA: principal component analysis
ROS: reactive oxygen species
SAH: S-adenosylhomocysteine
SAM: S-adenosylmethionine
ssGSEA: single-sample gene set enrichment analysis
SHP2: Src homology region 2 domain-containing phosphatase-2
TBHP: tert-butyl hydroperoxide
TIF: tumor interstitial fluid
OXPHOS: oxidative phosphorylation

## Data availability

RNAseq, scRNAseq, Metabolomics, Proteomics and all other data supporting the findings of this study are available from the corresponding author upon reasonable request.

## Acknowledgements

- Silke Hempel, Stephanie Mewes, Kerstin Meyer, Julia Kern and Antoine Devisme
- The authors thank the Core Facility AMIR^CF^ (DFG-RIsources N° RI_00052) for support in MR-imaging
- The authors thank the lighthouse Core Facility of the Medical Faculty, University of Freiburg (Project Numbers 2023/A2-Fol; 2021/B3-Fol), the DKTK, and the DFG (Project Number 450392965).
- The Proteomic Platform – Core Facility was supported by the Medical Faculty of the University of Freiburg to O.S. (2021/A3-Sch; 2023/A3-Sch).

