## Supplementary Figure 1 for "Vertical RAS-pathway inhibition in pancreatic cancer drives therapeutically exploitable mitochondrial alterations"

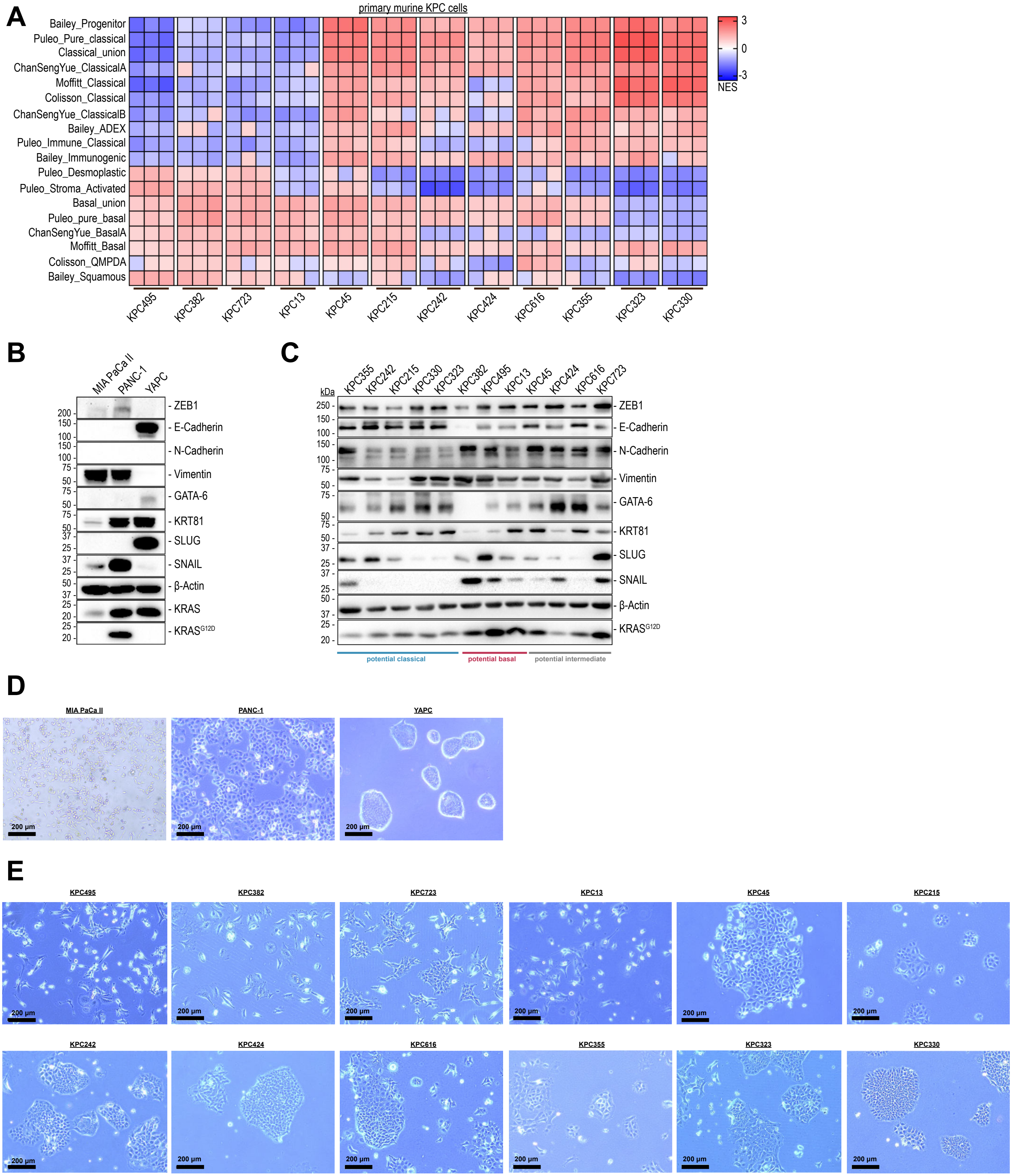

**Supplementary Figure 1: Molecular subtype classification of murine KPC cells and human PDAC cell lines.** (***A***) Transcriptomic analysis of murine KPC cells classifying them into basal-like or classical-like molecular subtypes. (***B***) Western Blot analysis of human PDAC cell lines for subtype related proteins. β-actin served as housekeeping protein. (***C***) Western Blot analysis of murine KPC cells for subtype related proteins. β-actin served as housekeeping protein. The protein lysates were generated between passage 5-10. (***D***) Morphology of human PDAC cell lines. (***E***) Morphology of murine KPC cells.
