## Supplementary Figure 2 for "Vertical RAS-pathway inhibition in pancreatic cancer drives therapeutically exploitable mitochondrial alterations"

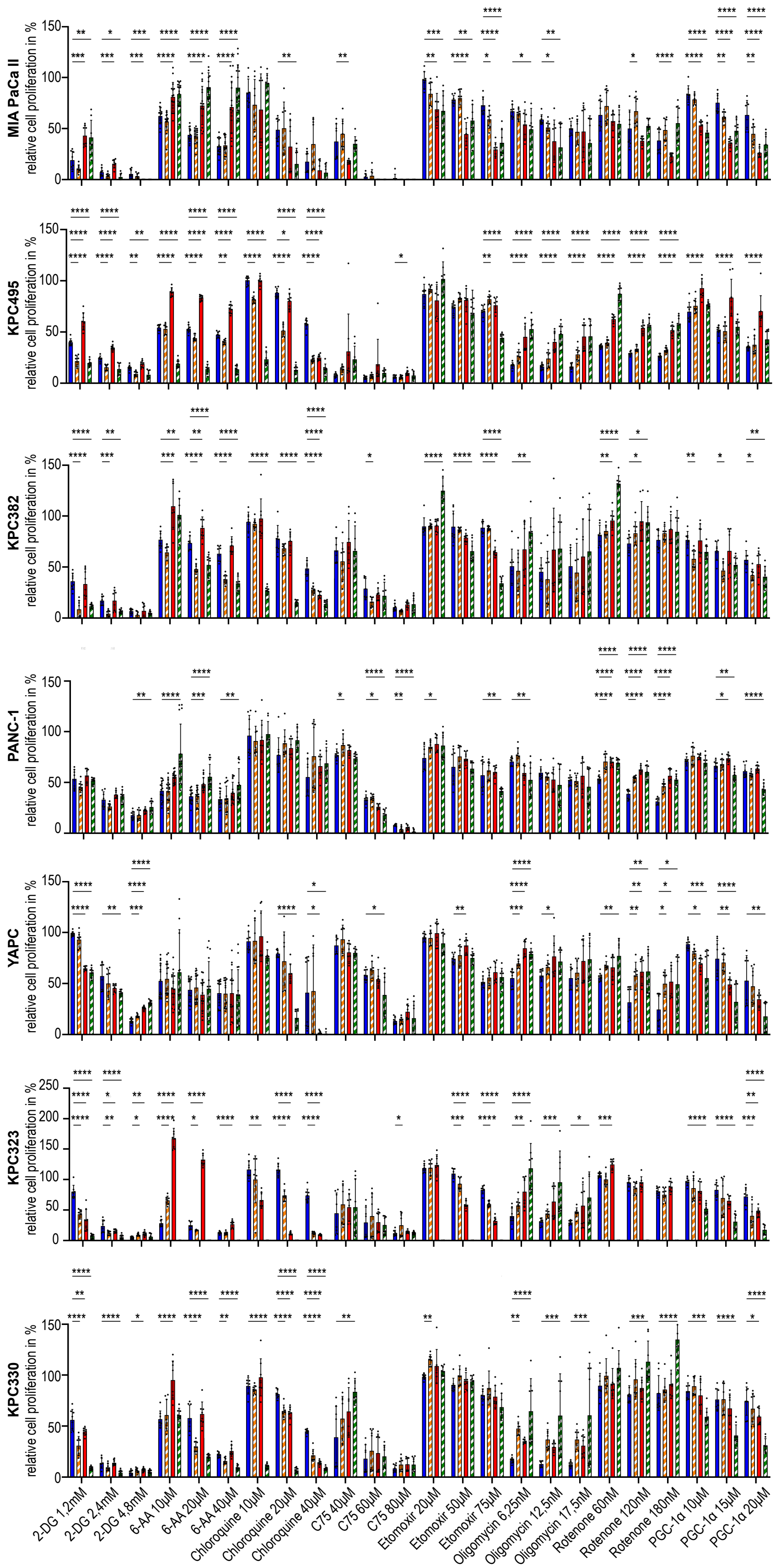

**Supplementary Figure 2: Influence of metabolic inhibition to SHP2 +/- MEK inhibition.** Proliferation assay of human PDAC cell lines and murine KPC cells. Cells were incubated for 5-8 days and treated with either DMSO, SHP099 (15 µM), Trametinib (10 nM), or combination of SHP099 and Trametinib. Additionally, the cells were treated with metabolic inhibitors at the depicted concentrations. To investigate the influence of the metabolic inhibitor on the related treatment option, the values are relative to their corresponding treatment without metabolic inhibition. Statistical significance was determined via one-way ANOVA with comparisons made only against corresponding DMSO controls.* $P < 0.05$, ** $P < 0.01$, *** $P < 0.001$, **** $P < 0.0001$
