## Supplementary Figure 3 for "Vertical RAS-pathway inhibition in pancreatic cancer drives therapeutically exploitable mitochondrial alterations"

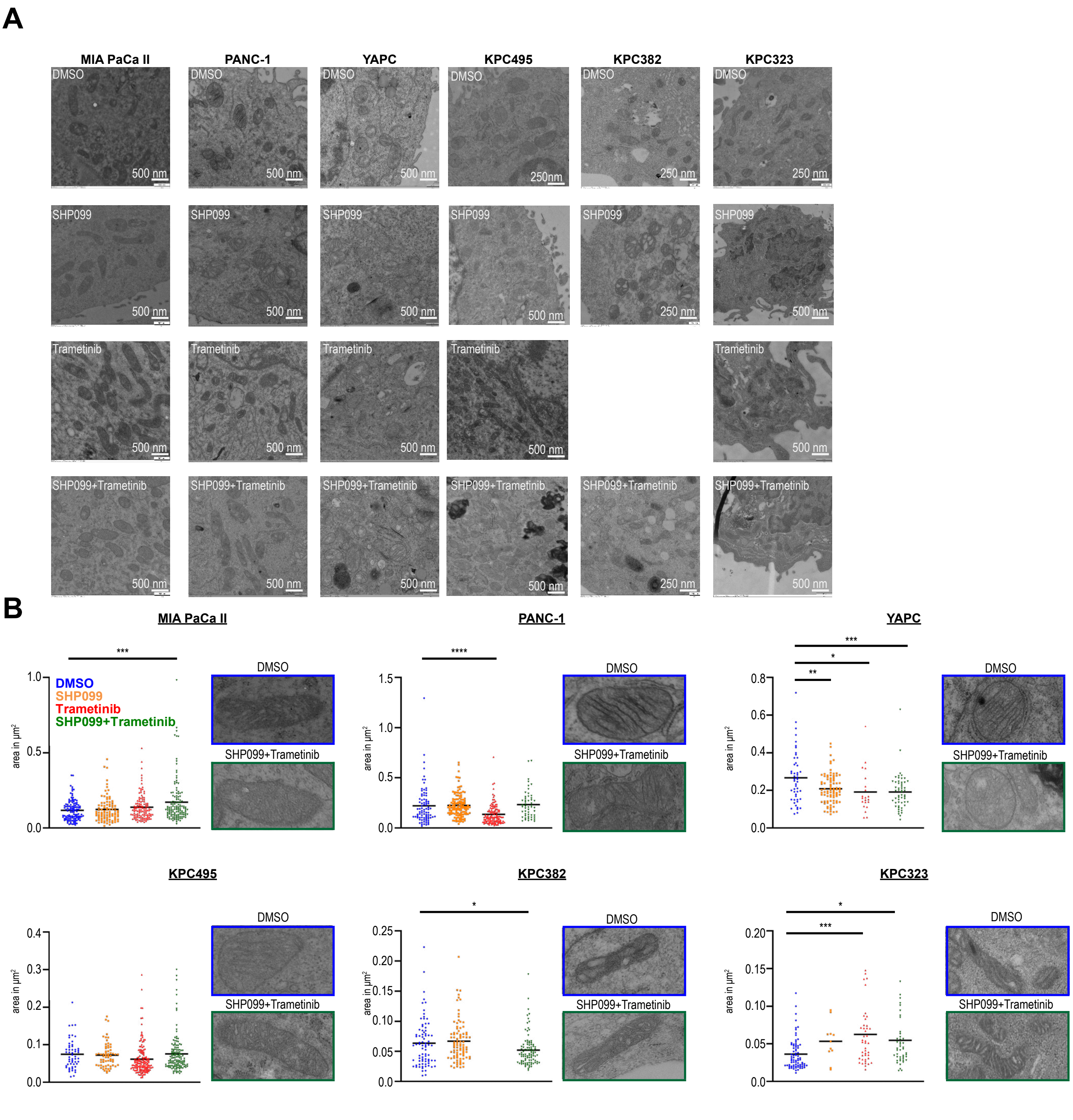

**Supplementary Figure 3: Dual inhibition of SHP2 and MEK initiates mitochondrial adaptions. (*A*)** Electron microscopy images of human PDAC cell lines and murine KPC cells. Cells were treated for 72 hours with either DMSO, SHP099 (15 µM), Trametinib (10 nM) or the combination of SHP099 and Trametinib. **(*B*)** Mitochondrial diameter of human PDAC cell lines and murine KPC cells based on electron microscopy. Statistical significance was assessed using one-way ANOVA in panel B with comparisons made against corresponding DMSO controls. * $P < 0.05$, ** $P < 0.01$, *** $P < 0.001$, **** $P < 0.0001$.
