## Supplementary Figure 4 for "Vertical RAS-pathway inhibition in pancreatic cancer drives therapeutically exploitable mitochondrial alterations"

# A

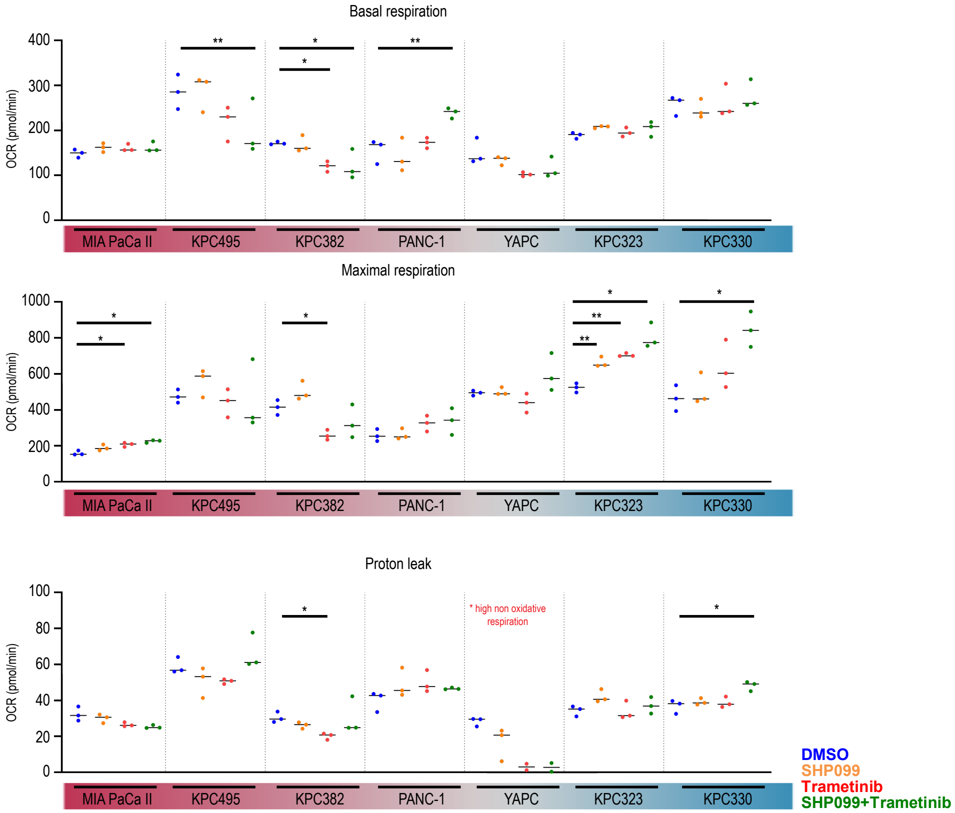


# B

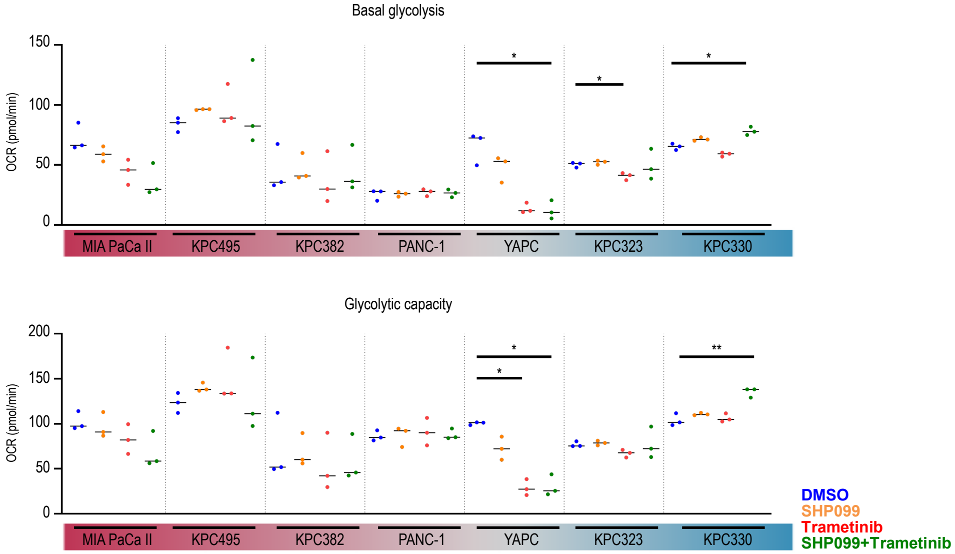

**Supplementary Figure 4: Dual inhibition of SHP2 and MEK leads to respiratory and glycolytic changes. (*A+B*)** Before measurement, human PDAC cell lines and murine KPC cells were treated for 72 hours with either DMSO, SHP099 (15 μM), Trametinib (10 nM) or the combination of SHP099 and Trametinib. **(*A*)** Respiratory behavior based on oxygen consumption rate. Mitochondrial function was assessed using the Agilent Seahorse XF Cell Mito Stress Test. **(*B*)** Glycolytic behavior based on extracellular acidification rate. Glycolytic function was measured using the Agilent Seahorse XF Glycolysis Stress Test. Data are shown as the mean of 3 independent experiments, with each biological replicate consisting of 6 technical replicates. Statistical significance was determined via one-way ANOVA in panels A and B, with comparisons made against corresponding DMSO controls.* $P < 0.05$, ** $P < 0.01$, *** $P < 0.001$, **** $P < 0.0001$
