## Supplementary Figure 5 for "Vertical RAS-pathway inhibition in pancreatic cancer drives therapeutically exploitable mitochondrial alterations"

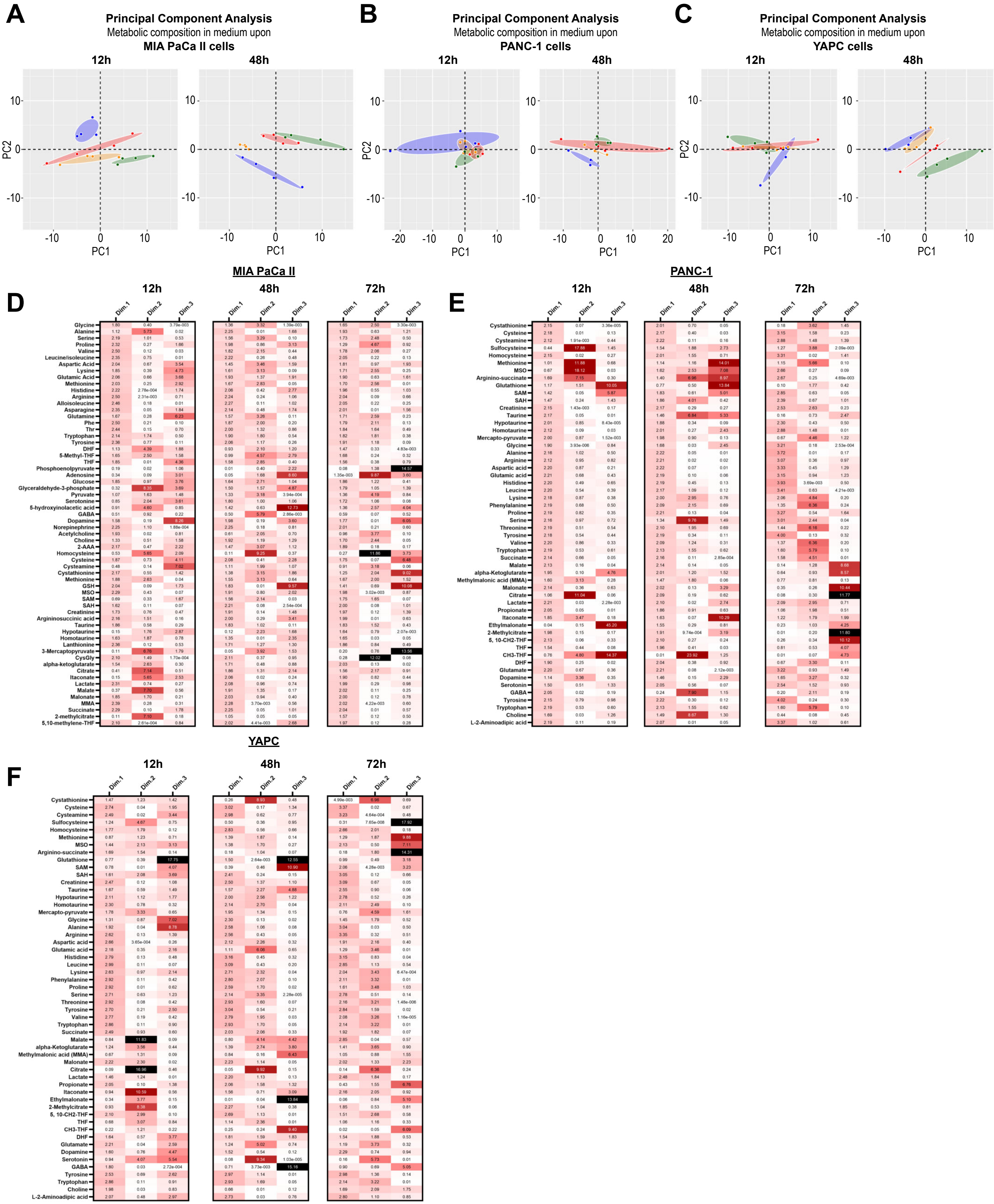

**Supplementary Figure 5: Impact of SHP2 +/- MEK inhibition on metabolite composition. (*A-C*)** Principal component analysis of extracellular metabolites from human PDAC cell lines after 12 hours and 48 hours of treatment with either DMSO, SHP099 (15 µM), Trametinib (10 nM) or the combination of SHP099 and Trametinib. Four independent samples are shown per treatment. Number of included metabolites: 62 for MIA PaCa II cells, 54 for PANC-1 and YAPC cells. **(*D-F*)** PCA contribution of human PDAC cell lines at 12, 48, and 72 hours of treatment. . The PCA contribution factors display the loadings of each variable on the principal components from the PCA. The contributions factors indicate how much each variable influences the principal components. Lower-dimensional principal components have a greater impact on the data, as they account for more of the variance.
