## Supplementary Figure 6 for "Vertical RAS-pathway inhibition in pancreatic cancer drives therapeutically exploitable mitochondrial alterations"

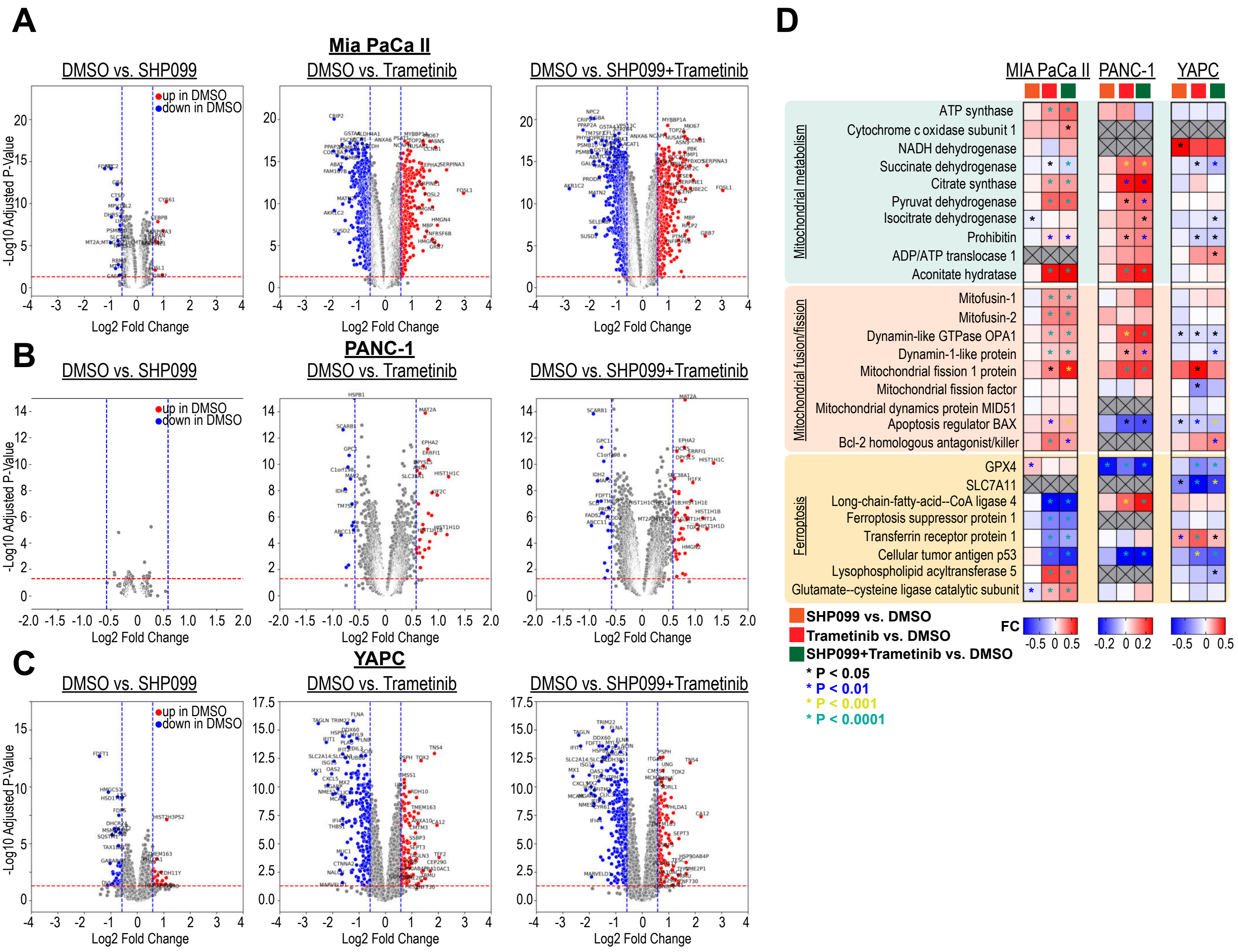

**Supplementary Figure 6: Proteomics analysis reveals enhanced mitochondrial metabolism.** (***A-C***) Proteomic profiling of human PDAC cell lines demonstrates altered protein abundance following 72 hours of treatment with SHP099 (15 μM), Trametinib (10 nM), or their combination, compared to control (DMSO). Displayed are proteins with Log2 fold changes < -0.58 or > 0.58, and adjusted P-values < 0.05. (***D***) Protein composition analysis reveals changes in mitochondrial metabolism, mitochondrial dynamics (fusion/fission), and ferroptotic response after 72 hours of treatment with SHP099, Trametinib, or their combination. Statistical significance for panels A-C was determined using Limma with a T-test and one-way ANOVA in panel D with comparisons made against corresponding DMSO controls. *P < 0.05, **P < 0.01, ***P < 0.001, ****P < 0.0001.
