## Supplementary Figure 7 for "Vertical RAS-pathway inhibition in pancreatic cancer drives therapeutically exploitable mitochondrial alterations"

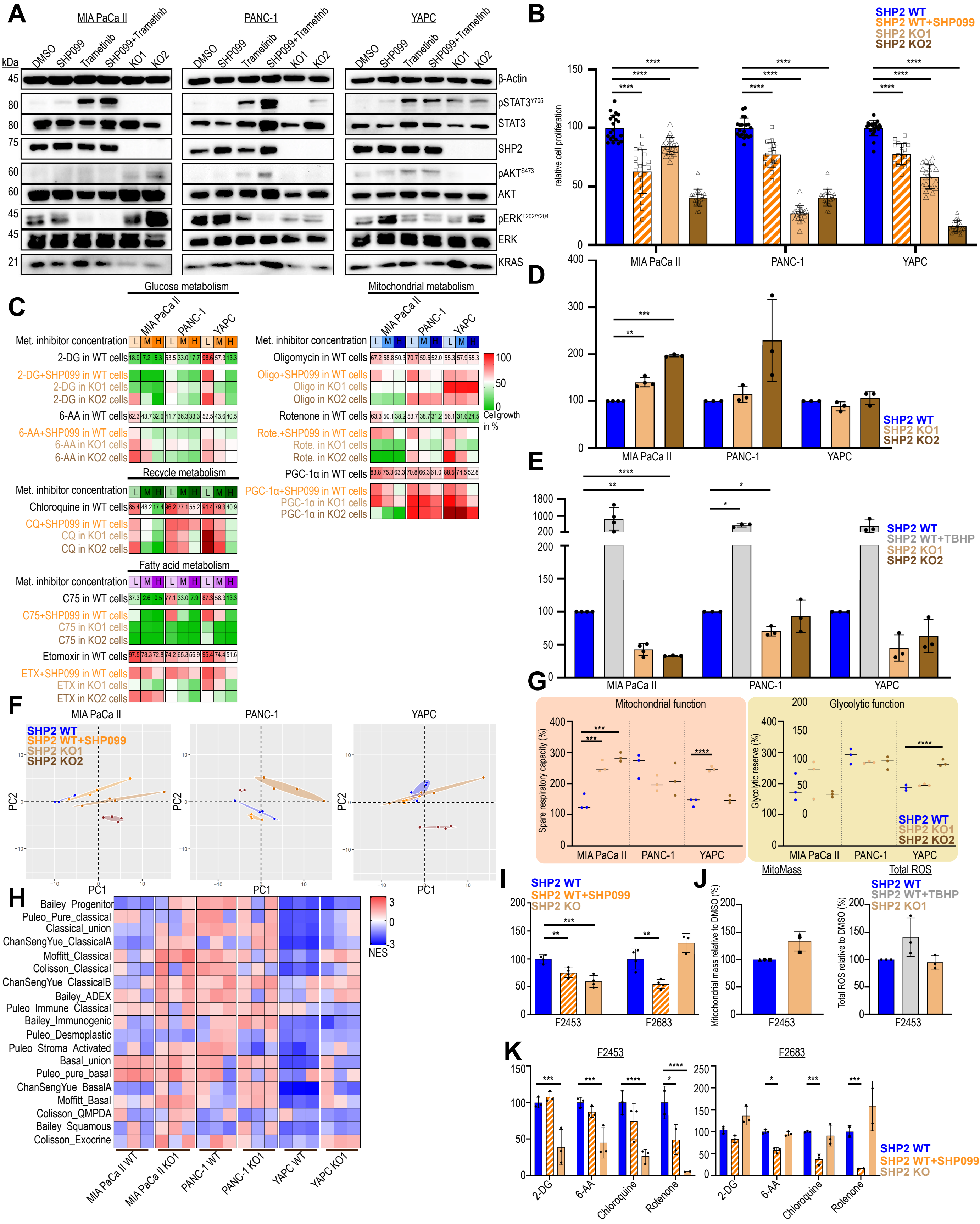

**Supplementary Figure 7: Genetic SHP2 knockout leads to molecular subtype switch and adapted metabolism. (*A*)** Western Blot analysis of human PDAC cell lines with and without genetic SHP2 KO. **(*B*)** Cell proliferation of SHP2 expressing cells and SHP2 KO cells for a period of 10 days. Error bars represent the SD of 20 wells from one experiment. The sensitivity to SHP2 treatment varied in this experiment compared to that depicted in Figure 1B. Here, the initial cell count was substantially lower, rendering them more sensitive to SHP099 in terms of proliferation compared to cells in Figure 1B. **(*C*)** Relative cell proliferation of SHP2 KO cells plus metabolic inhibition. The data for SHP2-expressing cells were extracted from the experiments shown in Fig. 1D-G. Each colored area of the heatmap represents the mean of a total of 8 wells from two independent experiments. **(*D*)** Flow cytometry determination of mitochondrial mass in human SHP2 expressing and KO cells relative to cell size. Each data point represents 1 of 3-4 independent experiments. **(*E*)** Flow cytometry determination of intracellular total reactive oxygen species in human SHP2 expressing and KO cells. Each data point represents 1 of 2-4 independent experiments. **(*F*)** PCA of human genetic SHP2 KO PDAC cell lines and SHP2 expressing PDAC cell lines upon 72h treatment. 4 independent samples are shown per treatment. Number of metabolites for MIA PaCa II cells: 62. Number of metabolites for PANC-1 and YAPC cells: 54. **(*G*)** Spare respiratory capacity and glycolytic reserve of human SHP2-expressing and SHP2 KO cells, based on ECAR and OCR values. Mitochondrial function was assessed using the Agilent Seahorse XF Cell Mito Stress Test and glycolytic function was measured using the Agilent Seahorse XF Glycolysis Stress Test. Data are shown as the mean of three independent experiments. **(*H*)** Transcriptomic analysis of SHP2 expressing and SHP2 KO cells indicate molecular subtype switches after genetic SHP2 loss. **(*I*)** Cell proliferation of primary murine PDAC cells. Each data point represents one independent experiment. **(*J*)** Flow cytometry determination of mitochondrial mass and total ROS in murine SHP2-expressing and SHP2 KO cells relative to cell size. Each data point represents 1 of 3 independent experiments. **(*K*)** Relative proliferation of SHP2 expressing cells treated with 15 µM SHP099 or SHP2 deleted cells in presence of metabolic inhibition. Values are relative to the respective DMSO treated control group. Statistical significance was determined via one-way ANOVA in panels B, D, E, G, I, J and K, with comparisons made only against corresponding DMSO controls.* $P < 0.05$, ** $P < 0.01$, *** $P < 0.001$, **** $P < 0.0001$
