## Supplementary Figure 8 for "Vertical RAS-pathway inhibition in pancreatic cancer drives therapeutically exploitable mitochondrial alterations"

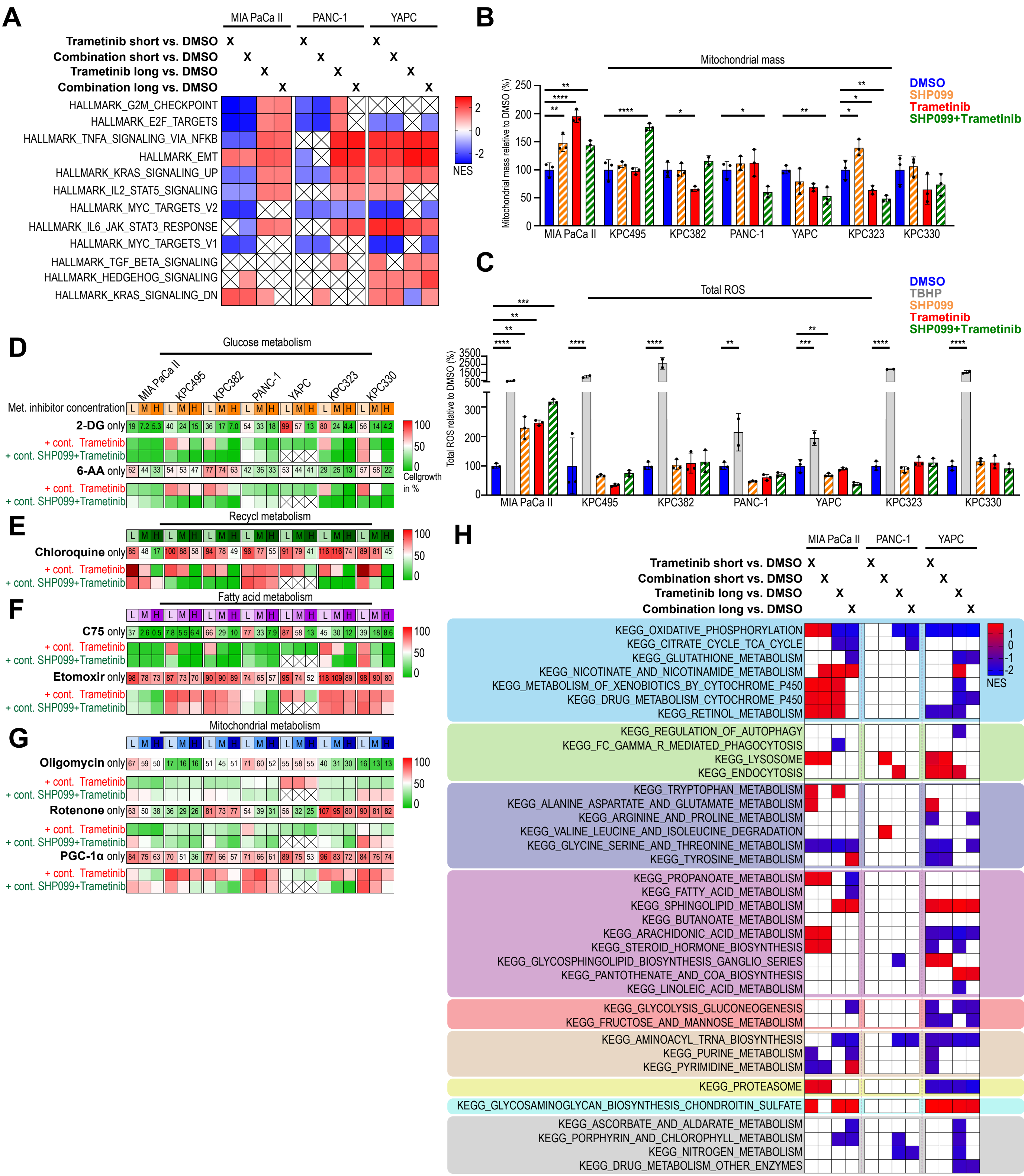

**Supplementary Fig. 8: Long-term MAPK inhibition leads to the development of resistance and sustained mitochondrial adaptations. A,** Transcriptomic analysis of human PDAC cell lines treated for 48 hours (short) or at least 8 weeks (long) with either Trametinib (10 nM) or a combination of SHP099 (15 µM) and Trametinib (10 nM). Gene set enrichment analysis (GSEA) was performed using Hallmark gene sets, with adjusted P-values < 0.25. The data represent treated cells relative to control cells. **B,** Flow cytometric analysis of mitochondrial mass in continuously treated human PDAC cell lines, normalized to cell size. Each data point represents 1 of 2-3 independent experiments. **C,** Flow cytometric analysis of intracellular total reactive oxygen species (ROS) in continuously treated human PDAC cell lines, normalized to cell size. Each data point represents 1 of 2-3 independent experiments. **D-G,** Relative cell proliferation in the presence of metabolic inhibitors (L: low, M: middle, H: high concentration) in potentially resistant human PDAC cell lines. Each colored area of the heatmap represents the mean of 8 wells from two independent experiments. Data for cells treated with only metabolic inhibitors were derived from Fig. 1 D-G. Metabolic inhibitor concentrations are similar to those from supplementary Fig. 2. The YAPC cells continuously treated with the combination therapy exhibited poor growth under these conditions and therefore could not be used in this approach. **H,** Gene set enrichment analysis of human PDAC cell lines treated for 48 hours (short) or at least 8 weeks (long) with either Trametinib (10 nM) or a combination of SHP099 (15 µM) and Trametinib (10 nM) based on the Kyoto Encyclopedia of Genes and Genomes (KEGG) pathways. Up- or downregulation is indicated by the corresponding squares, with gene sets having adjusted P-values < 0.25 highlighted. All white squares for these cell lines or treatment conditions are not statistically significant and are therefore excluded from the analysis.
