## Supplementary Figure 9 for "Vertical RAS-pathway inhibition in pancreatic cancer drives therapeutically exploitable mitochondrial alterations"

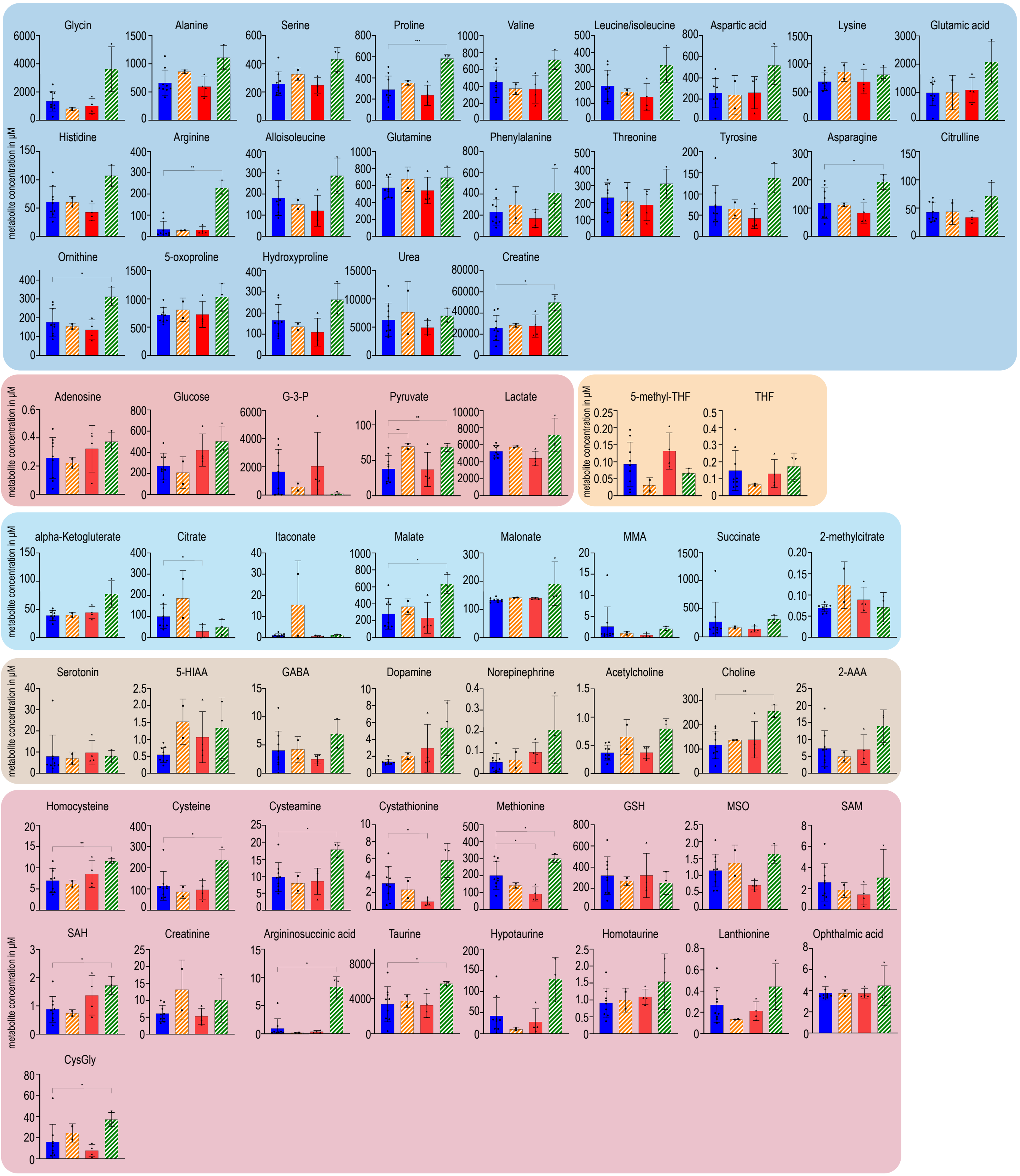

**Supplementary Figure 9: Pharmacological SHP2 and/or MEK inhibition influences tumor interstitial fluid metabolite composition.** Tumor interstitial fluids from freshly sacrified animals were utilized for measurement via LC-MS/MS. Animals were treated either with vehicle (acts as control), SHP099 (75mg/kg), Trametinib (1mg/kg) or the combination of SHP099 and trametinib. Statistical significance was determined via one-way ANOVA, with comparisons made only against corresponding DMSO controls.* P < 0.05, ** P < 0.01, *** P < 0.001, **** P < 0.0001.
