## Supplementary Figure 10 for "Vertical RAS-pathway inhibition in pancreatic cancer drives therapeutically exploitable mitochondrial alterations"

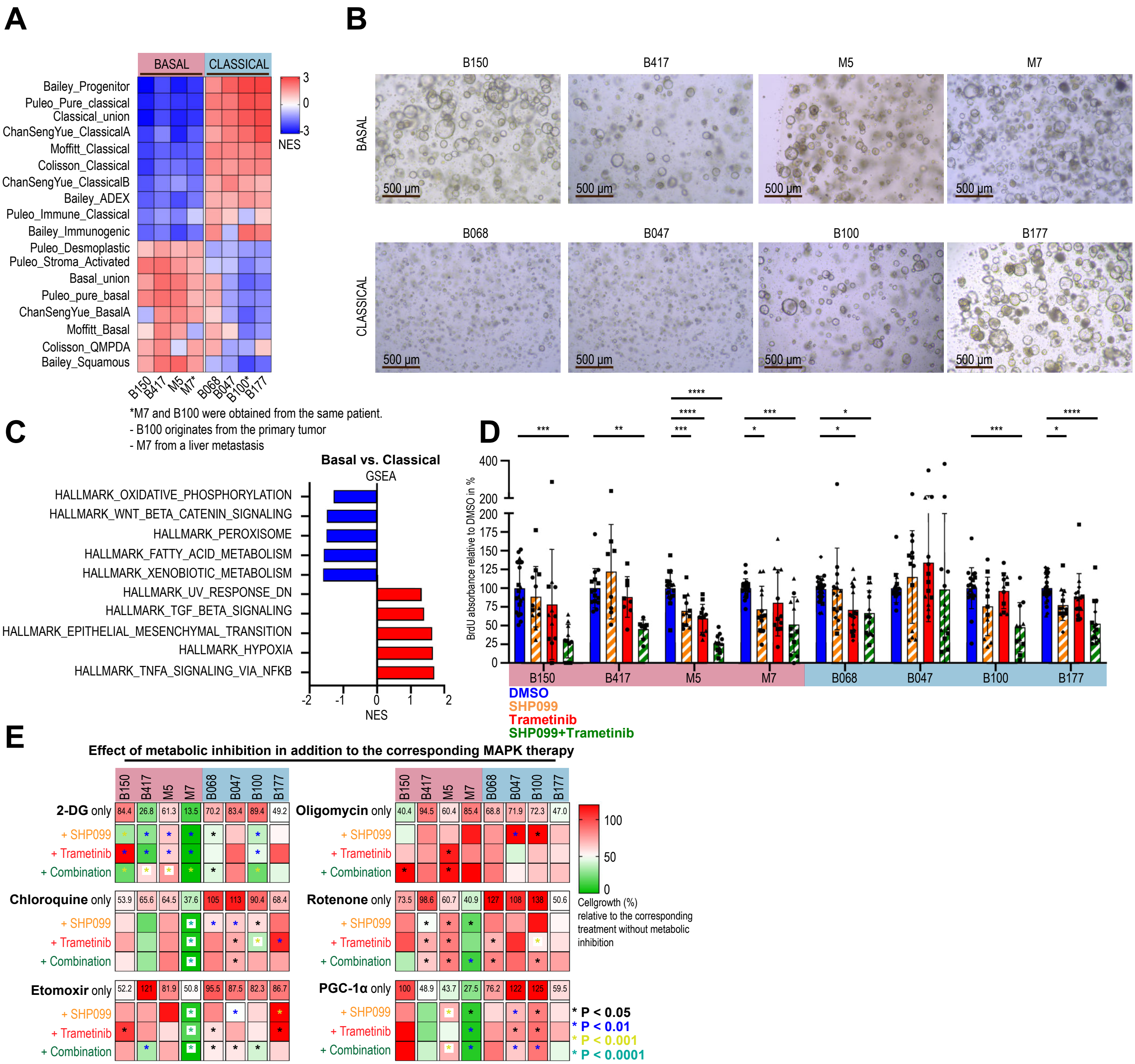

**Supplementary Figure 10: Mitochondrial adaptations in patient-derived PDAC organoids.**

(***A***) Transcriptomic analysis classifying human PDAC organoids into basal-like or classical-like subtypes. (***B***) Morphology of human-derived organoids. (***C***) GSEA comparing basal-like and classical subtypes using Hallmark gene sets. (***D***) Organoid proliferation in the presence of DMSO, SHP099 (15 μM), Trametinib (25 nM), or their combination, with 2-3 biological replicates (3-5 samples each). (***E***) Relative cell proliferation with or without MAPK inhibition plus metabolic inhibition. The data illustrate the effect of the metabolic inhibitor (e.g., 2-DG) on top of each MAPK inhibition. Data are normalized to MAPK inhibition without metabolic inhibitor (e.g., SHP099+2-DG vs. SHP099). Heatmap represents the mean of 4 samples. Metabolic inhibitor concentrations: 2-DG (2 mM), Chloroquine (20 μM), Etomoxir (50 μM), Oligomycin (30 μM), PGC-1αi (10 μM), Rotenone (20 nM). Statistical significance (D, E) was assessed via one-way ANOVA against DMSO controls: * $P < 0.05$, ** $P < 0.01$, *** $P < 0.001$, **** $P < 0.0001$
